# All-atom protein design via SE(3) flow matching with ProteinZen

**DOI:** 10.1101/2025.10.18.683228

**Authors:** Alex J. Li, Tanja Kortemme

## Abstract

The advent of generative models of protein structure has greatly accelerated our ability to perform *de novo* protein design, especially concerning design at coarser physical scales such as backbone generation and protein binder design. However, the design of precise placements at atomic scales remains a challenge for existing design methods. One avenue towards higher fidelity atomic-scale design is via generative models with full atomic resolution, but is complicated by the intricacies of simultaneously designing both discrete protein sequence and continuous atomic positionings. In this work we propose a framework to capture this interplay by decomposing residues into collections of oriented rigid bodies, allowing us to apply SE(3) flow-matching for all-atom protein structure generation. Our method, ProteinZen^1^, generates designs with high sequence-structure consistency while retaining competitive diversity and novelty on both unconditional and conditional generation tasks. We demonstrate competitive performance for unconditional monomer design and state-of-the-art performance on various forms of motif scaffolding, including full-atom motif scaffolding and motif scaffolding without specifying motif segment spacing or relative sequence order.

## 1 Introduction

*De novo* design has been greatly accelerated by generative modeling methods utilizing paradigms such as score-based diffusion [1] and flow matching [2], allowing us to design proteins with new structures *de novo* as well as simple functions such as protein-protein binders [3]. Despite these advances, targeting functions prevalent in nature, such as small molecule sensing or enzymatic catalysis, remains challenging. We posit this difficulty is in part due to the lack of atomic-level reasoning in existing generative models. The current dominant paradigm for protein design is backbone-centric: prominent models such as RFDiffusion [4] and Chroma [5] only model backbones, and extensions such as RFDiffusion2 [6] and LigandMPNN [7] introduce more atomic-awareness but still do not consider atomic structures in full detail. Moreover, pipelines utilizing these models often require generating 10^2^-10^3^ designs per suitable design candidate, of which only a fraction of candidates will ultimately have the function of interest after experimental validation. We know that atomic precision, both local and extended, is necessary for many design targets such as *de novo* enzymes [8][9] and allosteric regulation [10]. Hence, these shortcomings motivate transitioning towards all-atom generation, where all available atomic detail is considered and refined during the generation process.

The difficulty of transitioning from backbone generation to all-atom protein structures lies in the interdependencies between discrete and continuous aspects of protein structure: amino acid residues take discrete identities with discrete atomic compositions, but their associated atomic placements exist in continuous space. Hence, precise placements of atoms requires careful treatment of the interplay between these quantities. Various methods have proposed different residue representations, including superposition-based [11] [12] [13], factorized [14], hybrid physical-latent [15] [16], and purely latent representations [17] [18]. However, all-atom methods still lag behind backbone-centric approaches on unconditional generation benchmarks, especially when it comes to co-designing compatible sequences for generated backbones. Furthermore, existing methods have much lower *in silico* success rates on conditional design tasks (such as motif scaffolding tasks) when compared to unconditional generation tasks, motivating further development.

Inspired by the intuition of viewing molecules as collections of functional groups, in this work we propose decomposing protein residues into collections of atoms that form oriented rigid bodies. Our decomposition allows us to represent residues with three oriented rigid bodies while retaining full expressivity of any canonical amino acid. We then apply the framework of SE(3) flow-matching [19][20] to train ProteinZen, our generative model of all-atom protein structure using this representation. We demonstrate competitive performance on unconditional monomer generation and state-of-the-art performance on various motif scaffolding tasks, helping pave forward the direction of all-atom protein generative models.

## 2 Methods

### 2.1 Setup

In the following sections we largely focus on a single residue *X*^(*i*)^ at position *i*, with a protein being an ordered sequence of such residues. We will index fragments within residue *i* using local index *j* or globally using *k* (as a shorthand for *k* = (*i, j*)). Subscripts will be reserved for indexing along time/interpolation coordinate *t*. When clear we drop superscripts for simplicity, in which case it is implied that we are looking at a single entity (residue, residue fragment, etc.).

### 2.2 Representing all-atom detail using oriented rigid bodies

The residue representation in this work stems from the observation that every canonical protein residue can be described with at most three rigid chemical fragments (decomposition described in Appendix A). Hence, we can represent a residue as an ordered collection of three oriented rigid bodies *X* = (*S*, [*T*^(*a*)^, *T*^(*b*)^, *T*^(*c*)^]), where *S* ∈ 𝒜 is a residue identity in the twenty canonical residue identities 𝒜 and *T* = (*r, x*) ∈ SE(3), *r* ∈ SO(3), *x* ∈ ℝ^3^ is an oriented rigid body (also called a “frame”). We construct a mapping per residue identity between raw atomic coordinates and their frame-based representation (Appendix A, example in Figure 1A), which allows us to freely interconvert between the two representations. In this work, we train a generative model over rigid bodies and use this mapping to convert to and from cartesian atomic representations when necessary.

**Figure 1.**
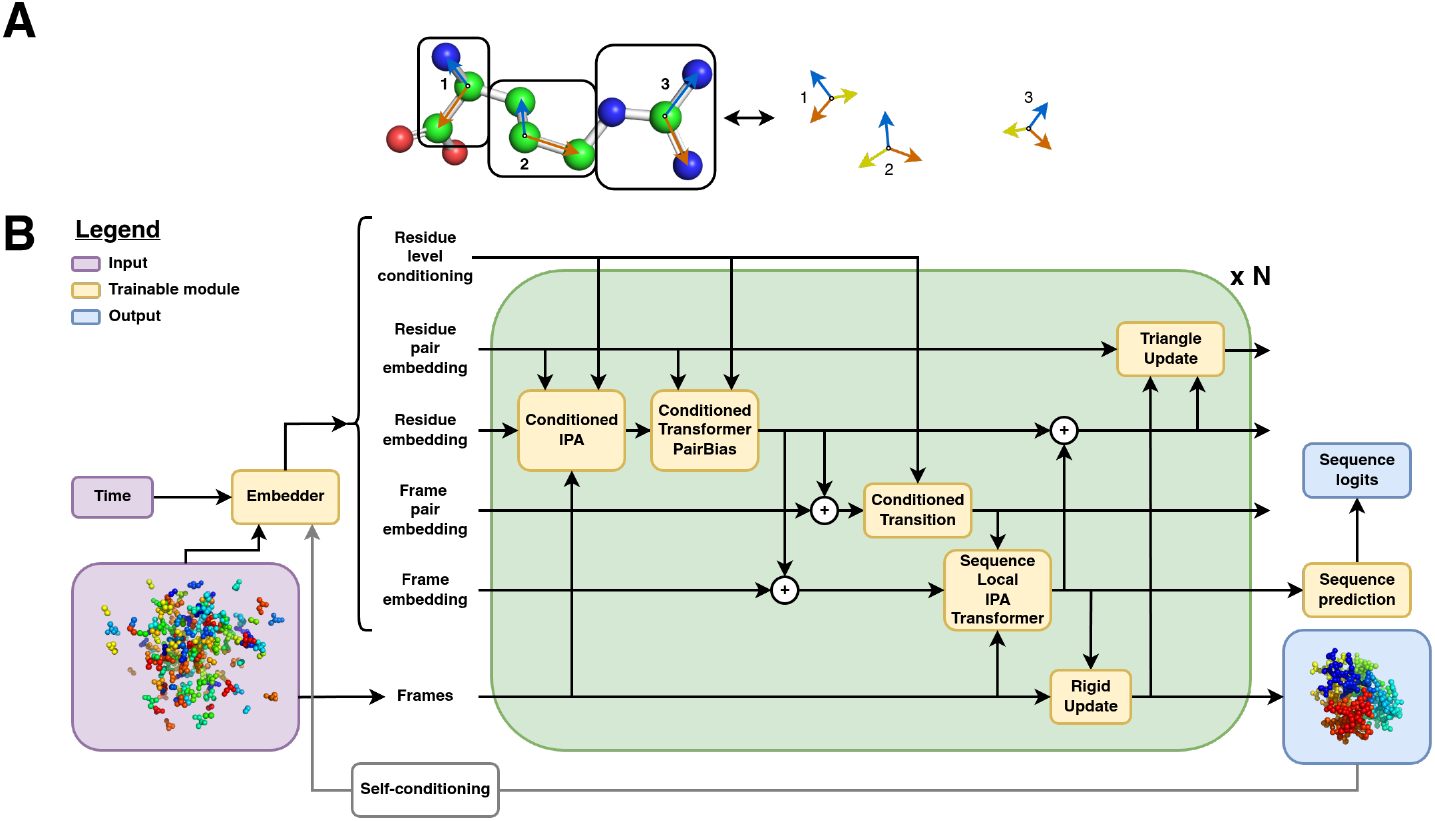
ProteinZen all-atom generation scheme. (A) Example decomposition of a protein residue into oriented rigid chemical groups. (B) ProteinZen architecture.

### 2.3 All-atom protein generation as SE(3) flow matching

We train our denoising model ProteinZen to minimize the SE(3) flow-matching objective over the collection of frames *{T*^(*k*)^ *}*. We follow FrameFlow [19] and FoldFlow [20] in flow-matching over SE(3) by training flows on SO(3) and ℝ^3^. More concretely, we construct the following interpolants from source/noise distribution *T*_0_ = (*r*_0_, *x*_0_) ~ *ρ*_0_ (see Appendix B) to target distribution *T*_1_ = (*r*_1_, *x*_1_) ~ *ρ*_1_ (the distribution of all-atom protein structures) along interpolation coordinate *t*:

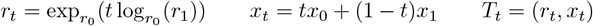

We formulate our flow-matching losses as denoising objectives and train a denoiser 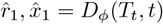 to minimize the following losses:

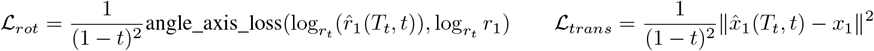

where ℒ_*rot*_ is a separable analog to the original SO(3) denoising objective. We further train a sequence prediction head with a cross-entropy loss ℒ_*seq*_ (which allows us to determine the sequence identity necessary for the atom-to-frame interconversion) and a FAFE-inspired auxiliary loss ℒ_*F AF E*_ [21] to improve global denoising properties (see Appendix D for further details).

In total, the full loss we optimize for is

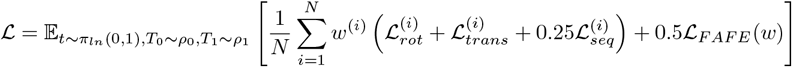

where *π*_*ln*_(0, 1) is the logit-normal distribution with location 0 and scale 1, and *w* specifies a residuewise loss weighting.

### 2.4 ProteinZen architecture

The architecture of ProteinZen shown in Figure 1B. We build a two-track architecture which extends IPA Transformers [22] to perform multi-scale modeling and also modify layers to allow training at scale (details in Appendix E). The use of frames and IPA as the primary units of computation makes ProteinZen fully SE(3) equivariant.

### 2.5 Sampling

Previous SE(3) flow-matching models sampled designs by integrating the following ODEs:

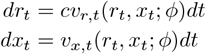

where *c* is a bespoke scaling of the rotation velocity field (often a large constant such as *c* = 10) at sample time which was found to be necessary for high quality sample generation [19] [20]. Although this trick can be used to generate high quality samples, it offers little tunability over the sampling process at inference time. Given that diffusion models in other literature have shown strong performance gains purely based on optimization of the integration strategy [23], and the space of samplers for SE(3) flow models remains underexplored, we aimed to create a better SE(3) sampling method.

Inspired by the connection between diffusion and flow matching models via stochastic interpolants [24] [25], we instead propose SDE sampling from an SE(3) flow model. We sample from ProteinZen by integrating the following SDEs:

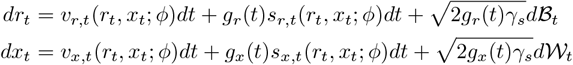

where *g*_*r*_(*t*), *g*_*x*_(*t*) control Langevin dynamics, *γ*_*s*_ controls the scale of noise added, and *v*_*x/r,t*_, *s*_*x/r,t*_ corresponded to the learned velocity and score functions, respectively. We estimate the velocity and score functions as follows:

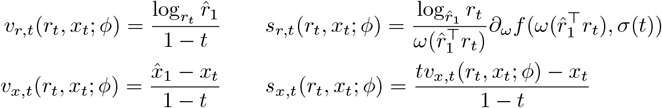

where 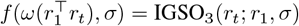 is the PDF of the IgSO(3) distribution centered at *r*_1_ and with variance *σ* and 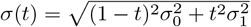 is the estimated variance of the rotation SDE. The ℝ^3^ score is derived from the connection between Gaussian flow-matching and diffusion processes [24] and the SO(3) score is computed as done in FrameDiff [22]. To our knowledge, this is the first example of SDE sampling from an SE(3) flow matching model trained on a deterministic flow. We also develop a custom integration strategy inspired by EDM’s stochastic integrator [23] which is further described in Appendix F.

## 3 Results

### 3.1 Unconditional Monomer Generation

#### 3.1.1 Metrics

We evaluate generation quality on three main criteria. The first is *sequence-structure consistency* (SSC), which we evaluate by predicting the structure of the generated sequence using ESMFold [27] and computing the all-atom RMSD (aaRMSD) against the generated structure. We use a threshold of aaRMSD < 2Å for a sample to be SSC. The second is *diversity* along both structure (DIV-str) and sequence (DIV-seq) axes, measured as the number of clusters formed when clustering with Foldseek [28] with a TM-threshold of 0.5 or MMSeqs2 [29] with default settings for structure and sequence, respectively. For both diversity metrics, we divide the number of clusters by the total number of SSC samples to yield the final metric. The third metric is *structural novelty* (NOV-str), where we query the PDB for the most structurally-similar hits using Foldseek and average the TM-scores of the maxTM hit per structure.

We also evaluate similar metrics for the backbone structures of generated samples. In this case, we swap the SSC metric for a designability metric (DES), where we use ProteinMPNN [30] to sample 8 sequences per backbone and predict structures for each using ESMFold. We define the minimum *C*_*α*_ RMSD between any of the predicted structures and the design structure as the self-consistent RMSD (scRMSD) and use a threshold of scRMSD < 2Å for a structure to be designable. The other two metrics DIV-str and NOV-str are evaluated on designable structures using the lowest scRMSD ESMFold prediction.

#### 3.1.2 Benchmark results

We benchmark ProteinZen and relevant comparison models on 100 samples for each length 70, 100, 200, and 300 residues (Table 1). ProteinZen has competitive performance in unconditional monomer generation, having a respectable 65% of samples being sequence-structure consistent, similar to Pallatom and only outperformed by concurrent work La-Proteina. We continue to recapitulate the observed trend that all-atom generation models generally have much lower SSC than DES, signaling a disconnect between generated backbones and their associated sequence and atomic detail. However, more recently developed models have began to close the performance gap.

**Table 1:**
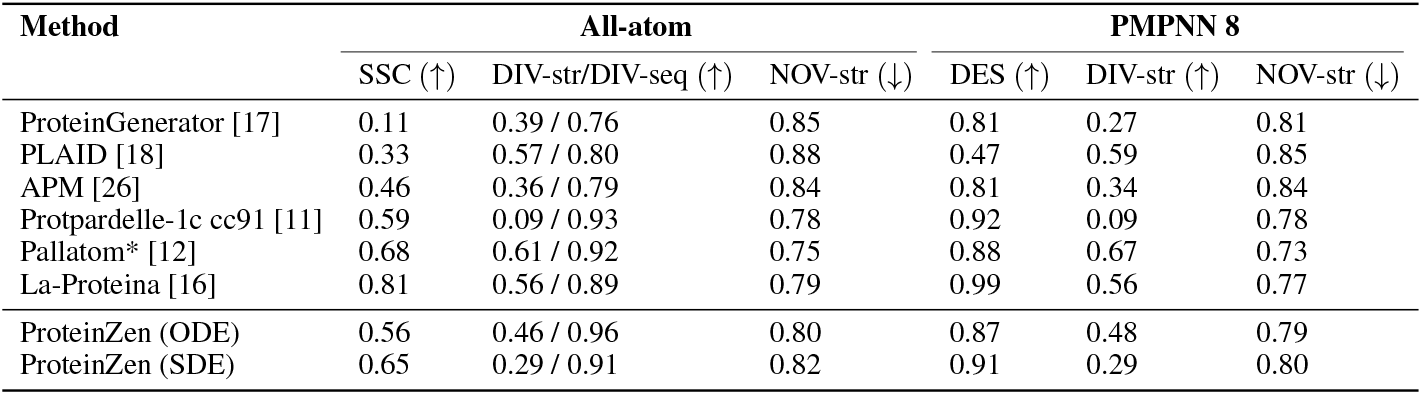
Sample quality metrics for ProteinZen and relevant comparison models. *Pallatom was trained on proteins of length at most 128, so the reported performance is partially out-of-distribution.

#### 3.1.3 SDE vs ODE sampling

In Table 1, we also evaluate the performance of ProteinZen using previously reported ODE samplers to provide a comparison point against our proposed SDE sampler. For the benchmark values reported, we use the SDE sampler with a sampling parameter set we found to maximize SSC, yielding a significant performance boost on SSC at the cost of structural diversity. However, we find the SDE sampler also achieves a better tradeoff between diversity and SSC than using the previous ODE method and can be controlled by tuning the noise scale parameter *γ*_*s*_ (Figure F.1).

#### 3.1.4 Sample characteristics

We evaluate other characteristics of ProteinZen samples in Figure 2. Figure 2A shows SSC as a function of sequence length. We note that ProteinZen performance drops off as protein length increases: this may be explained by the increased difficulty of designing larger proteins because the space of possible protein structures increases as sequence length increases, and making it more challenging to ensure the designed sequence specifies the design structure.

**Figure 2.**
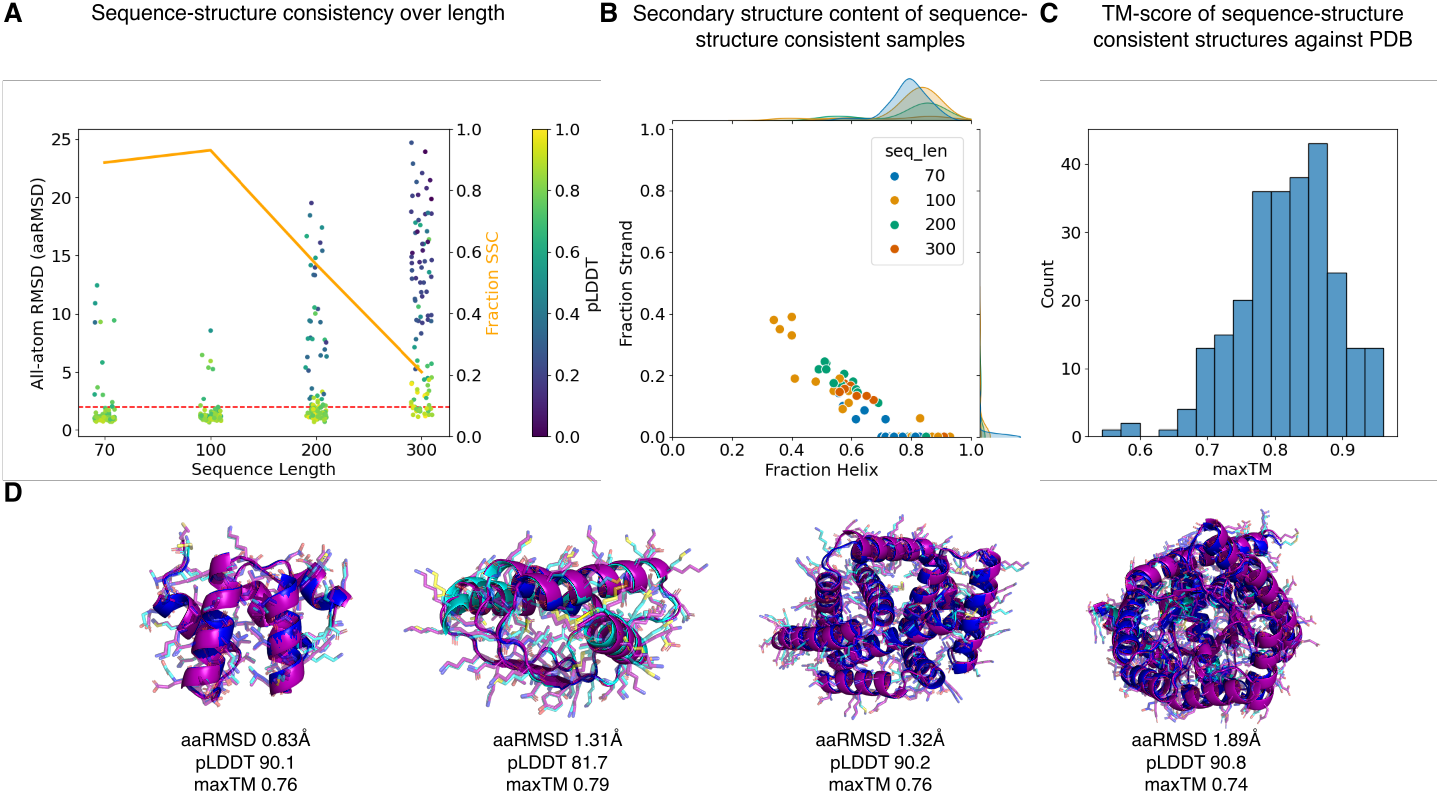
Various methods of evaluating ProteinZen sample characteristics. (A) Sequence-structure consistency of all-atom structures over length. (B) Secondary structure content of sequence-structure consistent samples. (C) maxTM scores of SSC samples against PDB. (D) Examples of sequence-structure consistent samples. Purple is the generated structure and blue is the ESMFold prediction colored by pLDDT.

Figure 2B shows the secondary structure distribution of SSC samples, depicting that overall the model is capable of generating structures with diverse secondary structures composition. When compared to the training data distribution (Appendix C), we observe that samples are biased towards *α*-helices in content, a property observed in many existing generative models [4][5]. We note that *β*-sheets have much higher contact order (involve contacts which are local in space but distal in sequence) than *α*-helices, and the design of such structures is an area of interest for further investigation.

Figure 2C depicts the maxTM distribution across SSC samples. All samples have maxTM > 0.5, signaling that ProteinZen has learned and largely recapitulates the known fold space of the PDB. There may be hints of generalization to novel fold space given the left tail end of lower maxTM scores (structures in Figure G.1), and improving this generalization is an avenue of future work.

Finally, Figure 2D shows examples of sequence-structure consistent samples. We generally get good agreement between the generated structure and the predicted model, with the core more well constrained than the surface as expected. We note that we occasionally observe minor unphysicalities, such as clashes or overextended bond lengths, but that these occur in a small minority of samples and may be addressed by finetuning with physical constraints (e.g. clash losses or bond length losses) or inference-time steering [31].

### 3.2 Motif scaffolding

Although methods have continually improved at unconditional generation tasks, motif scaffolding has remained a more challenging task. We evaluate motif scaffolding on the Protpardelle benchmark set [11] under two scenarios: (1) one-shot all-atom motif scaffolding and (2) motif scaffolding with sequence redesign. For models with such capabilities, we also evaluate performance on both *indexed motif scaffolding* (motif fragment order and spacing is pre-specified) and *unindexed motif scaffolding* (must infer both motif fragment order and spacing).

#### 3.2.1 Metrics

Our primary metric is *unique success rate* per task, which can intuitively be understood as the number of high quality unique scaffolds generated per set of design attempts. For one-shot motif scaffolding, we define a particular design to be a “success” for a particular task if the global heavy atom RMSD < 2Å, the motif heavy atom RMSD < 2Å, and the motif C*α* RMSD < 1Å between the design structure and structure of the generated sequence predicted using ESMFold [27]. For models which do not impute the input all-atom motif structure, we further filter generated outputs by ensuring that there is no larger than 2Å heavy-atom RMSD pairwise between the motif structure in the design, the motif structure in the ESMFold prediction, and the input motif specification. To compute the unique success rate per task, we cluster task successes with Foldseek [28] with TM-threshold 0.5 and normalize by the total number of designs generated.

For motif scaffolding with sequence redesign, we use ProteinMPNN to redesign 8 sequences for the backbone scaffold (the motif sequence stays fixed). We then replace the global heavy atom RMSD criteria with a global backbone RMSD < 2Å threshold. When comparing against RFDiffusion [4], we graft the input motif structure into the proper location before computing any relevant metrics.

#### 3.2.2 One-shot all-atom motif scaffolding

One-shot all-atom motif scaffolding serves to evaluate the performance of all-atom generative models against one another. We compare ProteinZen against concurrently developed models La-Proteina [16] and Protpardelle-1c [13], where La-Proteina can be run in both indexed and unindexed mode but Protpardelle-1c can only be run in indexed mode. In indexed mode (Figure 3A) ProteinZen outperforms concurrently developed methods on unique success rate, solving 25/26 tasks and performing the best at 18/26 of them. In unindexed mode (Figure 3B) ProteinZen and La-Proteina have more comparable performance, with ProteinZen solving 24/26 tasks and performing the best at 12/26 of them. We show examples of successful indexed motif scaffolding designs in Appendix Figure H.1 and unindexed motif scaffolding designs in Appendix Figure H.2.

**Figure 3.**
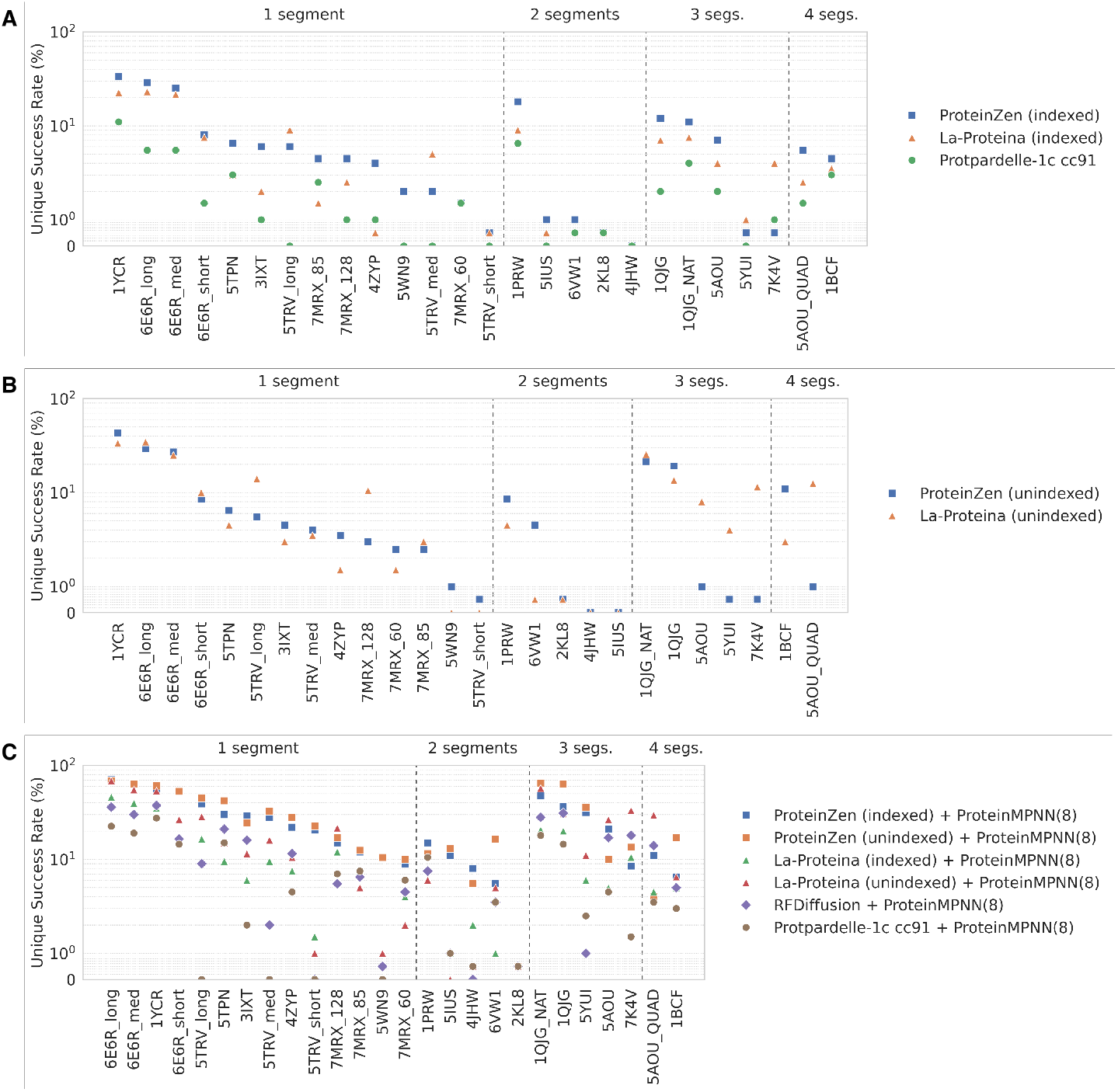
Performance on Protpardelle motif scaffolding benchmark. (A) Indexed one-shot all-atom motif scaffolding (B) Unindexed one-shot all-atom motif scaffolding (C) Motif scaffolding with sequence redesign.

#### 3.2.3 Motif scaffolding with sequence redesign

To evaluate how all-atom models compare against existing methods, we benchmark against RFDiffusion [4] which is currently the most popular model for protein motif scaffolding. To enable fair comparison, we evaluate motif scaffolding using ProteinMPNN [30] to redesign the sequences of generated scaffolds (the motif sequence remains fixed). Results are shown in Figure 3C. We find that ProteinZen consistently yields the best performance when compared against all models, solving all 26 tasks and performing the best on 21/26 tasks. Notably, the unique task success rate of all all-atom models is significantly improved by sequence redesign, highlighting that ensuring consistency between co-designed sequence and generated backbone is crucial to performant motif scaffolding as well.

## 4 Conclusions

We present ProteinZen, a method which utilizes flow-matching over rigid oriented chemical groups to generate all-atom protein structures. It exhibits promising performance on all-atom generation tasks, where 65% of unconditional monomer samples are sequence-structure consistent with atomic accuracy, and also yields state-of-the-art performance on the majority of benchmarked motif scaffolding tasks. Future work will extend ProteinZen to various other conditional generation tasks, including the binding of nucleic acids, small molecule, and protein targets.

## Acknowledgments and Disclosure of Funding

We would like to thank Dru Myerscough, Tobias Dorer, Hunter Nisonoff, Ishan Gaur, and Junhao Xiong for helpful discussions. A.J.L. is in part supported by a National Science Foundation Graduate Research Fellowship (NSF GRFP) and the UCSF Discovery Fellows program. T.K. is a Chan Zuckerberg Investigator. Portions of this work were performed on the Wynton HPC Co-Op cluster, which is supported by UCSF research faculty and UCSF institutional funds. The authors wish to thank the UCSF Wynton team for their ongoing technical support of the Wynton environment.

## Appendix

### A Residue frame construction

In this work, we define the decomposition of residues into rigid chemical fragments. To make the decomposition more unified, we use a common specification of the backbone fragment as

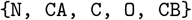

where `CB` is included for all residues that contain it (e.g. not `GLY`). We construct the frame as done in AlphaFold2 [32] from the `[N, CA, C]` atoms and follow FrameFlow [19] in computing the `O` coordinates from ideal peptide bond geometries. We then decompose the residue sidechain as follows (in a method inspired by that used in EquiFold [33]):

~~~
ALA: none
ARG: [NE, NH1, NH2, CZ], [CB, CG, CD]
ASN: [CG, ND2, OD1]
ASP: [CG, OD1, OD2]
CYS: [CA, CB, SG]
GLN: [CG, CD, NE2, OE1]
GLU: [CG, CD, OE1, OE2]
GLY: none
HIS: [CG, CD2, ND1, CE1, NE2]
ILE: [CB, CG1, CD1], [CB, CG1, CG2]
LEU: [CG, CD1, CD2]
LYS: [CD, CE, NZ], [CB, CG, CD]
MET: [CG, SD, CE]
PHE: [CG, CD1, CD2, CE1, CE2, CZ]
PRO: [CB, CG, CD]
SER: [CA, CB, OG]
THR: [CB, CG2, OG1]
TRP: [CG, CD1, CD2, CE2, CE3, NE1, CH2, CZ2, CZ3]
TYR: [CG, CD1, CD2, CE1, CE2, OH, CZ]
VAL: [CB, CG1, CG2]
~~~

For each rigid group, we define a set of axes to that allows us to construct the frame from atomic coordinates:

~~~
ALA: none
ARG: [NH1, CZ, NH2], [CB, CG, CD] ASN: [CG, ND2, OD1]
ASP: [OD1, CG, OD2]
CYS: [CA, CB, SG]
GLN: [CD, NE2, OE1]
GLU: [OE1, CD, OE2]
GLY: none
HIS: [ND1, CE1, NE2]
ILE: [CB, CG1, CD1], [CG2, CB, CG1] LEU: [CD1, CG, CD2]
LYS: [CD, CE, NZ], [CB, CG, CD] MET: [CG, SD, CE]
PHE: [CE1, CZ, CE2]
PRO: [CB, CG, CD]
SER: [CA, CB, OG]
THR: [CB, OG1, CG2] TRP: [CH2, CZ2, CZ3] TYR: [CE2, OH, CE1]
VAL: [CB, CG1, CG2]
~~~

where in a list [*x, y, z*] we construct the frame using Gram-Schmidt orthonormalization from vectors (coord(*x*) − coord(*y*)) and (coord(*z*) − coord(*y*)).

In order to create a representation expressive enough for all the canonical amino acids, we need to ensure each residue is represented with three oriented rigid bodies. For residue types with fewer than three rigid components, we supplement the representation with the frame defined by the first sidechain rigid group listed above. If there are no sidechain frames, the backbone frame is duplicated twice instead.

### B Noise distribution

Following FrameFlow [19], during training we use data-dependent coupling to sample from our noise distribution. For the rotation noise, we sample from the IgSO3 distribution centered around the ground truth data and with *σ* = 1.5, which helps push noise samples away from areas where we get degenerate solutions for the geodesic. For the translation noise, we sample from a standard isotropic Gaussian with *σ* = 1.6nm and align this noise against the ground truth data using the Kabsch algorithm.

During sampling we sample from the uniform distribution over SO(3) for rotations and an isotropic Gaussian with standard deviation *σ* = 1.6nm.

### C Training dataset construction

We curate and train ProteinZen on a custom dataset which we call AFDB512-clusters. We construct AFDB512-clusters using two different data sources: the PDB [34] and the AlphaFold predicted structures database [35]. For PDB structures, we adapt the strategy from FrameDiff [22] and select structures which meet the following criteria:

- X-ray structure or EM structure with resolution < 3.0 Å.
- Oligomeric state is specified as “monomer” in the mmCIF header.
- Length is between 31 and 512 residues.
- Greater than 50% of residues are in defined secondary structure elements as defined by DSSP [36]

We cluster these structures based on the 40% sequence identity cluster assignments specified by the PDB. This results in 22,518 structures total.

For AFDB structures, we adapt strategies from Protpardelle [11] and Pallatom [12] and select structures which meet the following criteria:

- pLDDT > 80.
- Length is between 31 and 512 residues.
- Greater than 50% of residues are in defined secondary structure elements as defined by DSSP [36].
- Longest loop < 15 residues.
- More than 2 secondary structure elements as defined by DSSP.
- Structures are >30% core, where a core residue is defined as a residue with at least 18 neighboring *C*_*α*_ atoms within 10 Å.
- Radius of gyration is less than 30 Å.

We first apply these filtering criteria to all AFDB Foldseek cluster representatives [37], then to all cluster members for the retained clusters. This results in 2,522,042 structures total.

**Figure C.1:**
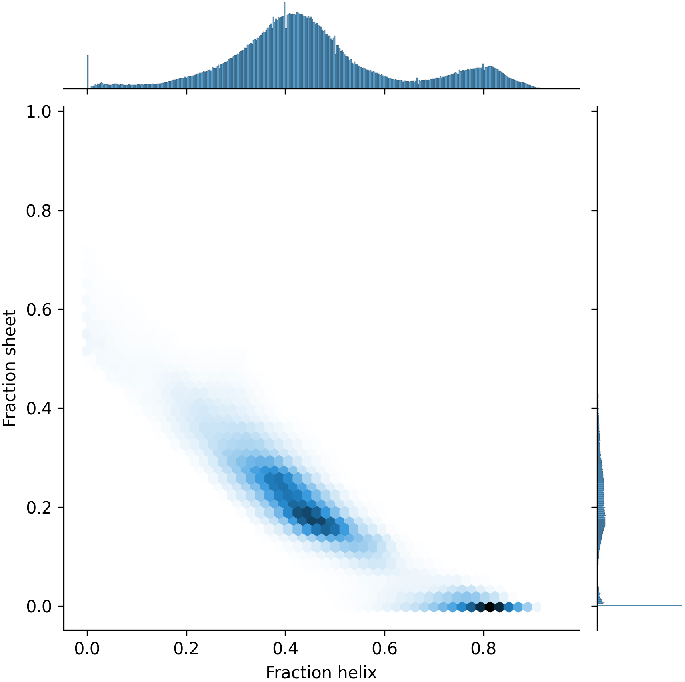
AFDB512-clusters secondary structure composition

### D Training details

As mentioned in the main text, we formulate our flow-matching losses as denoising objectives and train a denoiser 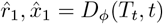 to minimize the following losses:

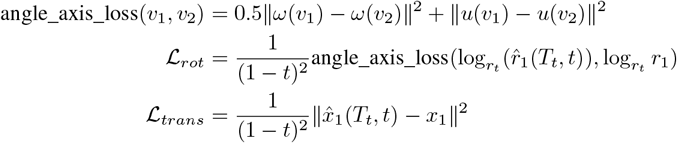

where *ω*(*v*) is the length of *v* and *u*(*v*) is the normalized vector of *v*. This form of ℒ_*rot*_ is a separable analog to the original SO(3) denoising objective inspired by FrameDiff [22] which we found led to more stable training in early experiments. As our frame-to-atom mapping requires a residue identity as input, we train a small prediction head at the end of the denoising network via cross-entropy loss to predict the ground-truth sequence, which allows us to decode out the identity of the residue at the end of a denoising trajectory:

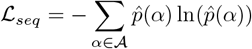

Finally, inspired by Wu et al. [21], we add an auxiliary loss ℒ_*F AF E*_ to encourage better global denoising properties via pairwise denoising terms.

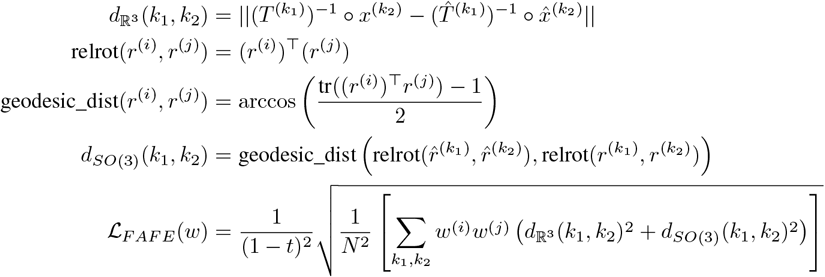

In total, the full loss we optimize for is

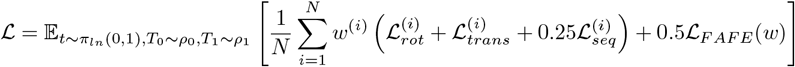

where *π*_*ln*_(0, 1) is the logit-normal distribution with location 0 and scale 1, and *w* is a residue-wise loss weighting term.

In practice we find better performance when using a residue-identity-based loss upweighting for *w*, and we set

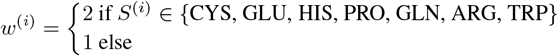

We suspect that this improves performance because it reduces the tendency of the model to excessive design alanines under reduced-noise sampling schemes.

#### D.1 Pretraining

We perform pretraining in multiple stages as listed in Table D.1. We follow Genie2 [38] in generating motif scaffolding training tasks but cap the max motif size to either 50% of the protein structure or 20 residues, whichever is smaller. Additionally, we select the denoising task for each motif residue independently as follows:

- Specify the entire residue with probabiliy 0.30. The residue sequence identity is provided.
- Specify only the backbone frame of the residue with probability 0.35. The residue sequence identity is masked.
- Specify only the tip frame of the residue with probability 0.35. The residue sequence identity is provided.

We note that pretraining was performed in one shot, and further tuning the training regime and hyperparameters could yield performance benefits and reduce training time. Training was performed on 8 NVIDIA H100s GPUs.

**Table D.1:**
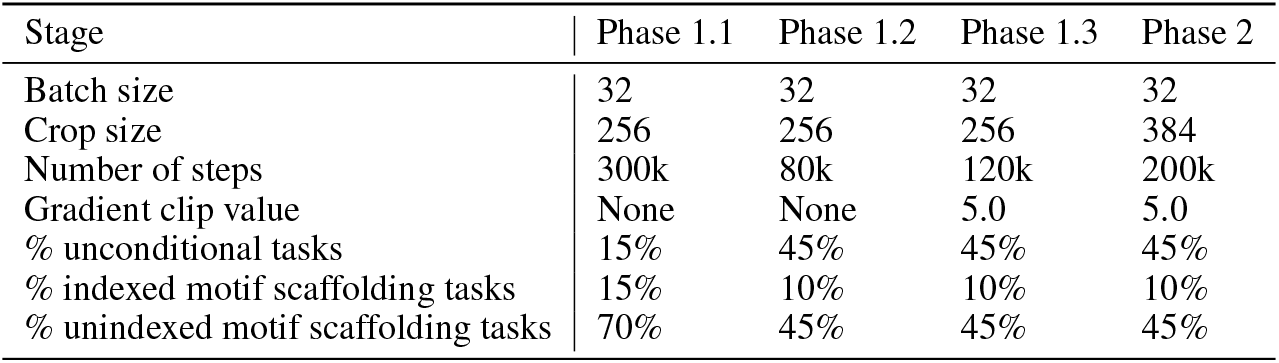
Pretraining regime.

| Stage | Phase 1.1 | Phase 1.2 | Phase 1.3 | Phase 2 |
| --- | --- | --- | --- | --- |
| Batch size | 32 | 32 | 32 | 32 |
| Crop size | 256 | 256 | 256 | 384 |
| Number of steps | 300k | 80k | 120k | 200k |
| Gradient clip value | None | None | 5.0 | 5.0 |
| % unconditional tasks | 15% | 45% | 45% | 45% |
| % indexed motif scaffolding tasks | 15% | 10% | 10% | 10% |
| % unindexed motif scaffolding tasks | 70% | 45% | 45% | 45% |

We sample from both PDB and AFDB data such that the ratio of real-to-synthetic structures is 1:3, and sample structures with weight inversely proportional to the structure’s associated cluster size

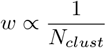

We use the Adam optimizer with *β*_1_ = 0.9, *β*_2_ = 0.999 and a base learning rate of 0.0001. After 300k steps, we start to decay the learning rate by 0.95 every 20k steps.

#### D.2 Motif scaffolding finetuning

From the Phase 2 checkpoint, we train a motif scaffolding specific set of weights by further finetuning on motif scaffolding as described in Table D.2. We warm start the learning rate back to 1e-4 and allow it to decay as before every 20k steps. We also change the max motif size to be either 50% of the protein structure or 160 residues, whichever is smaller.

**Table D.2:** Motif scaffolding finetuning regime.

| Stage | FT-Motif |
| --- | --- |
| Batch size | 48 |
| Crop size | 320 |
| Number of steps | 80k |
| Gradient clip value | 5.0 |
| % indexed motif scaffolding tasks | 20% |
| % unindexed motif scaffolding tasks | 80% |

### E ProteinZen architecture details

#### E.1 Invariant Point Attention Variants

In order to improve performance and computational efficiency while retaining the benefits of the original algorithm, we make several variants of Invariant Point Attention (IPA) which incorporate different adjustments.

##### Sequence-local IPA

Inspired by AlphaFold 3 [39], we perform efficient local modeling by restricting attention to pre-defined sequence-local blocks. In this work, per sequence-local block, we use 16 query frames and 64 key frames.

##### Pre-LayerNorm IPA

Invariant Point Attention in its original formulation followed earlier trends using post-layer norm in Transformer-style blocks. In line with modern Transformers, we instead switch to using pre-layer norm IPA.

##### Ada-LN IPA and output gating

Given pre-layernorm IPA, we also create a variant that enables conditioning via adaptive layer normalization techniques such as AdaLN [40] and output gating.

##### *QK-norm IPA* Vanilla

Transformers have found that normalization of the query and key vectors prior to attention calculations helps improve training stability [41]. We found this to be the case for IPA as well, and implement QK-normalization using a learnable LayerNorm for both query and key values.

#### E.2 Rotation preconditioning

Previous IPA-based SE(3) flow-matching models note an explosion in the learned rotation velocity norm as *t* → 1 which results in poor sample quality, necessitating a tampering of the velocity at inference-time at times close to *t* = 1 using some scaling function, commonly a multiplicative factor of (1 − *t*). We choose to directly embed this scaling factor into the model itself by scaling the magnitude of denoising prediction by (1 − *t*). We call this technique “rotation preconditioning” as inspired by EDM preconditioning [23], and note that is numerically equivalent to the model regressing against the rotation velocity field directly rather than the denoising target. Although rotation preconditioning did not yield any significant performance differences during small-scale experiments, it allows us to simplify the interpretation and manipulation of the velocity field output at sample time.

#### E.3 Motif input specification

Rather than impute and freeze the target motif as input specification, we instead append the motif as additional residues/frames and specify these tokens are motif inputs. In the indexed motif scaffolding case, we provide sequence indices as input, but in the unindexed case sequence indices are masked. During training, no motif imputation is performed. At inference time, at the end of a denoising trajectory, the model will compute which residues correspond to motif residues and imputes the motif at these residue positions.

#### E.4 Algorithm descriptions

**Figure E.1:**
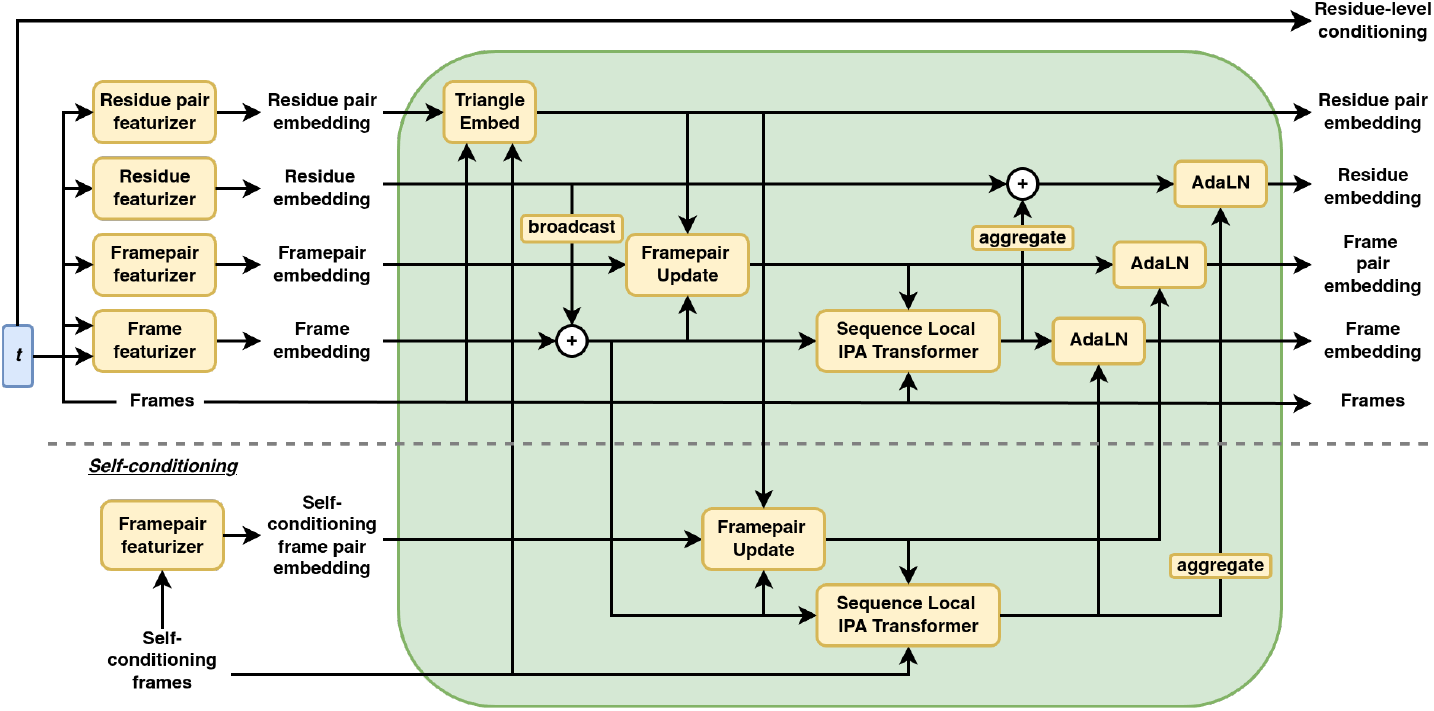
ProteinZen embedder architecture

We sketch pseudocode for some notable algorithms in ProteinZen (both the embedder in Figure E.1 and the denoiser in Figure 1B) in the algorithms below. We use *i, j* to index residue-level features and *l, m* to index frame level features. We further distinguish between residue- and frame-level input features by adding hats to all frame-level input quantities. Official implementations can be found at https://github.com/alexjli/proteinzen.

##### Algorithm 1

residue_featurizer

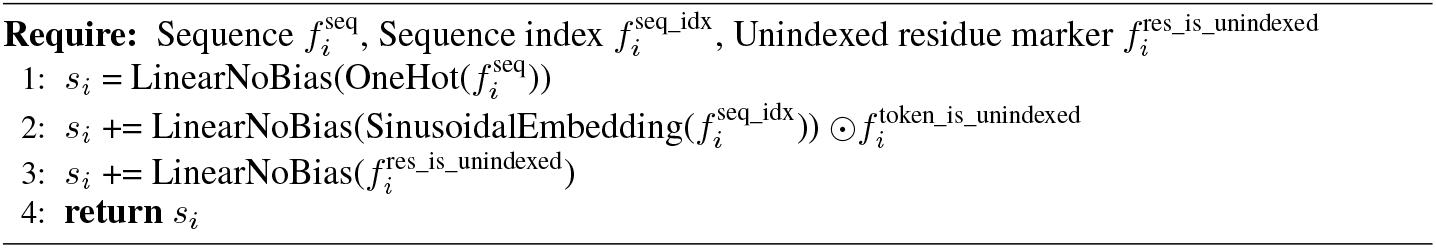

##### Algorithm 2

residue_pair_featurizer

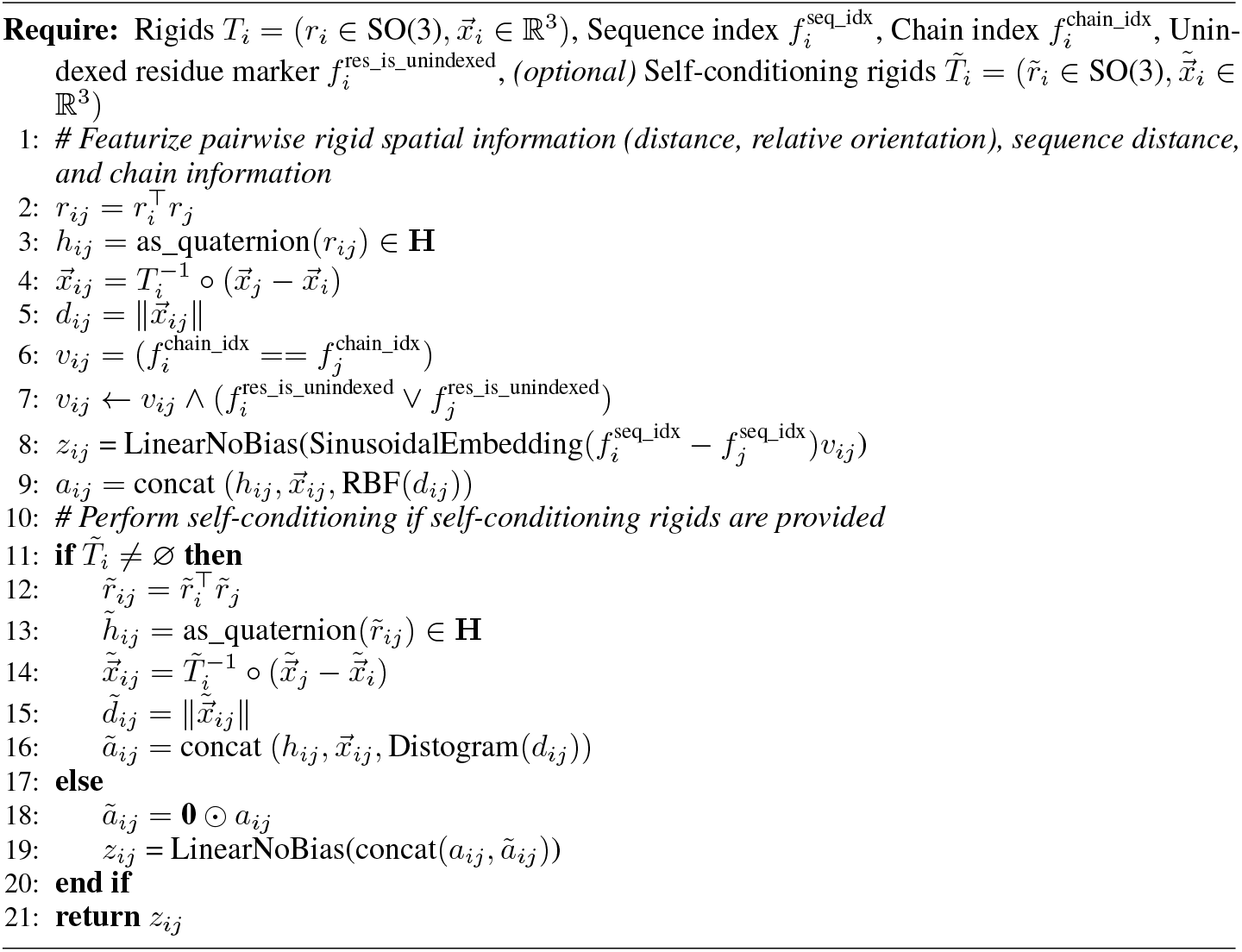

##### Algorithm 3

frame_featurizer

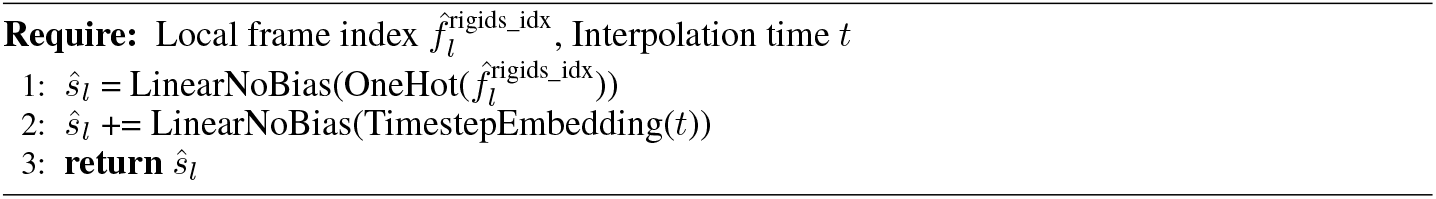

##### Algorithm 4

framepair_featurizer

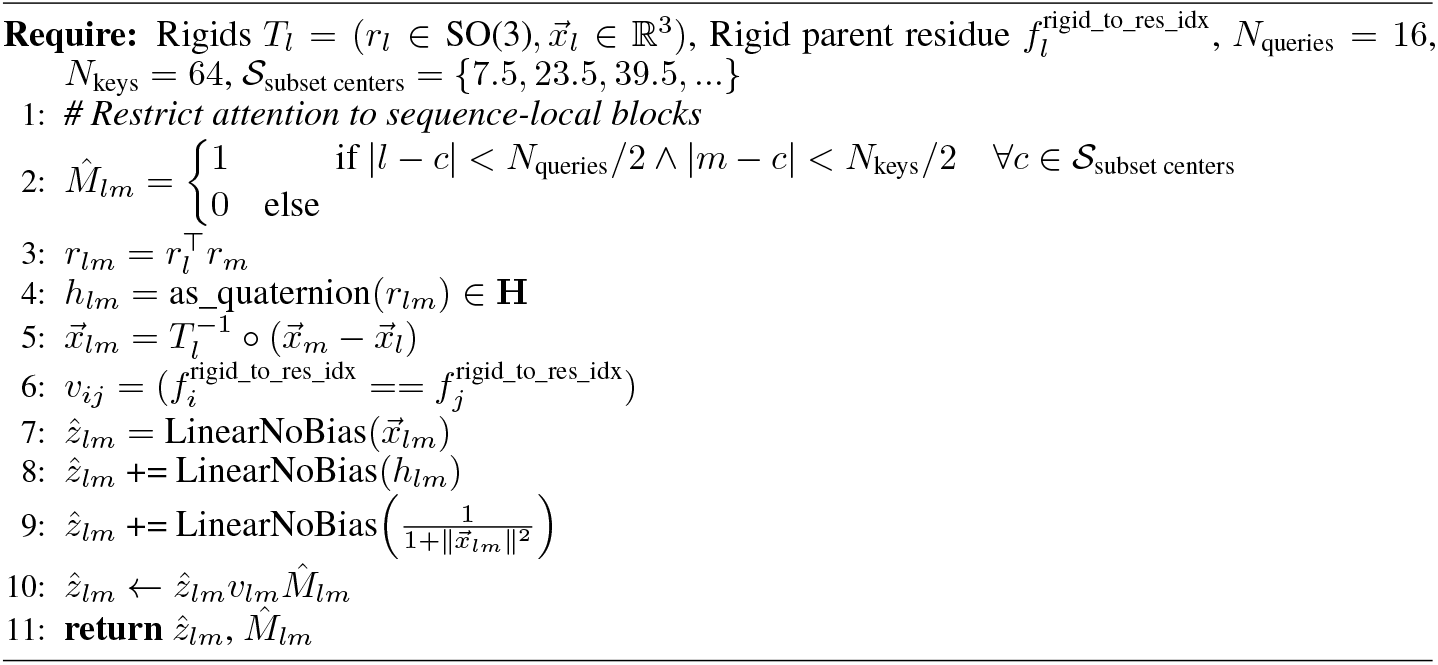

##### Algorithm 5

triangle_embed

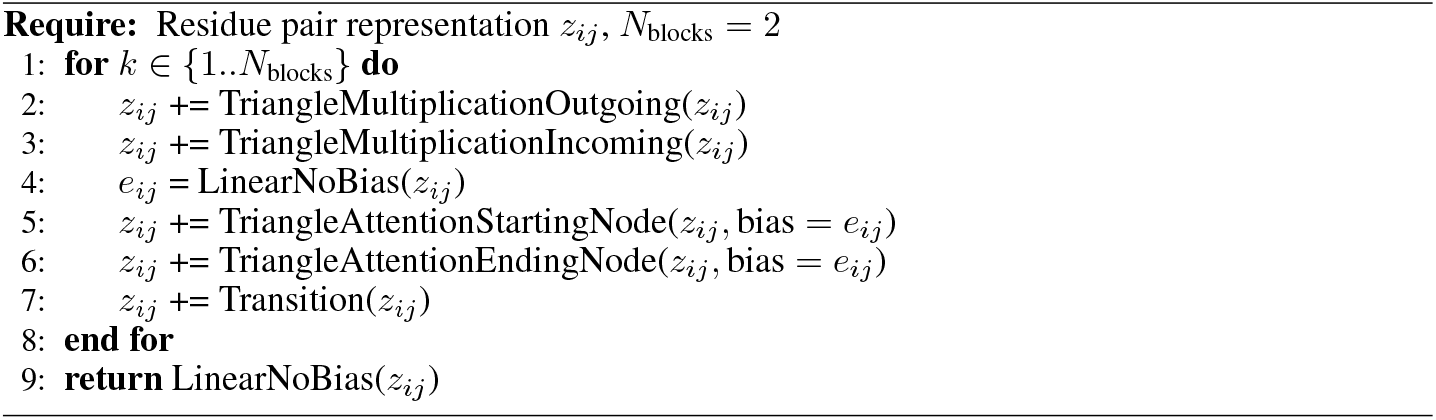

##### Algorithm 6

framepair_update

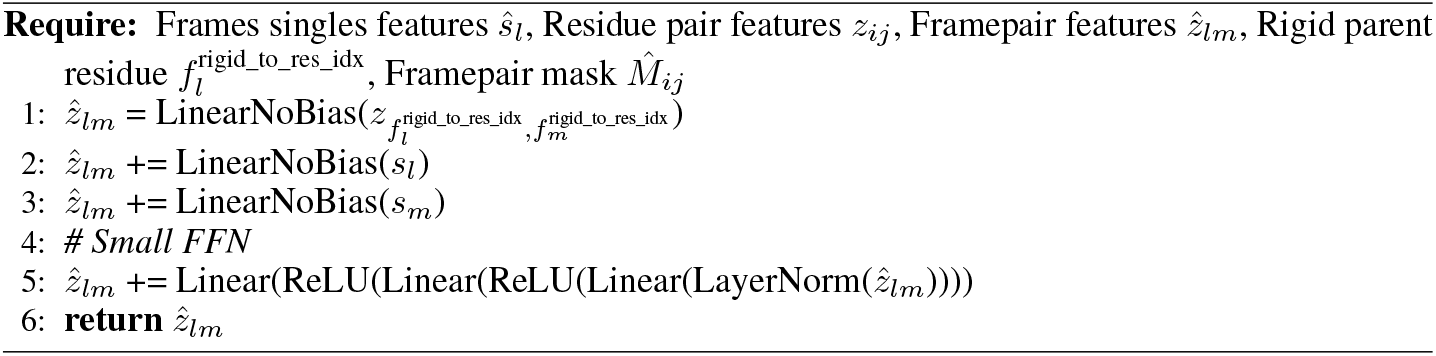

##### Algorithm 7

conditioned_ipa

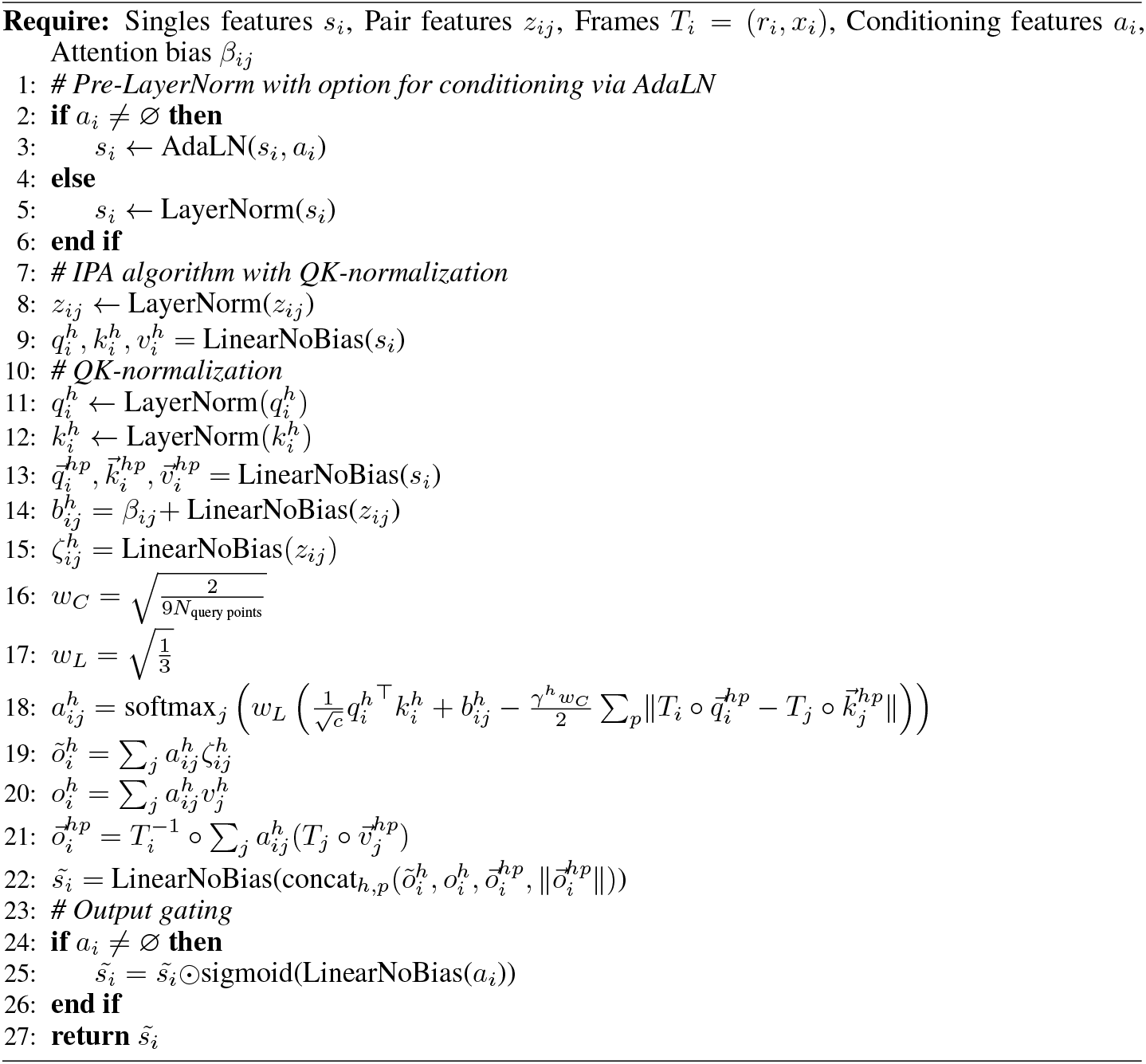

##### Algorithm 8

sequence_local_ipa

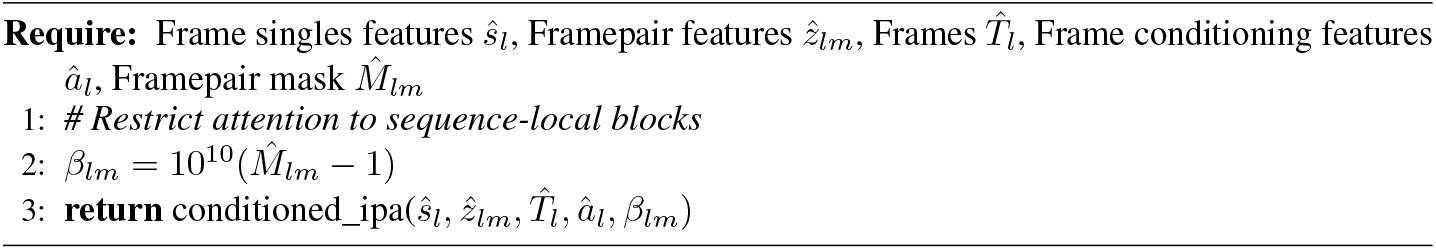

##### Algorithm 9

sequence_local_ipa_transformer

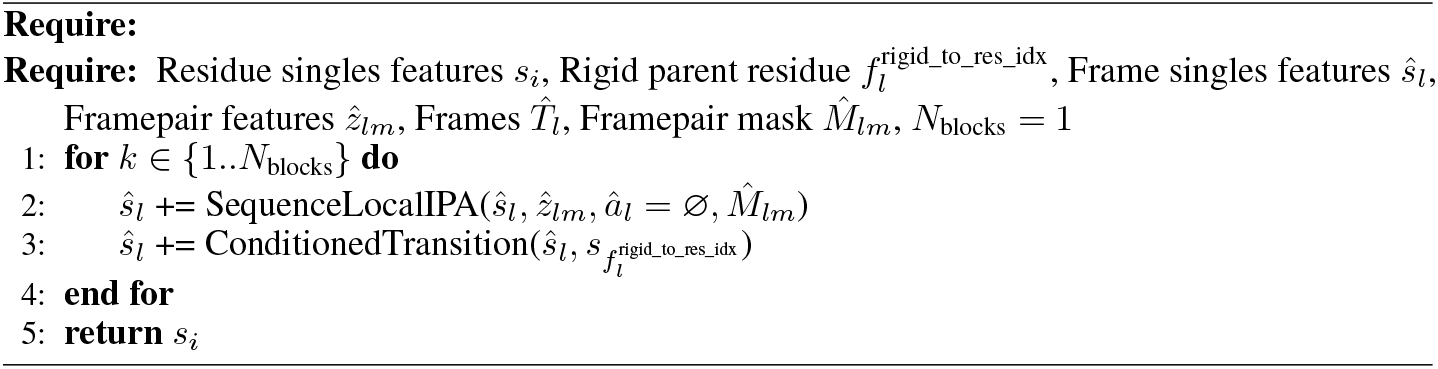

##### Algorithm 10

conditioned_self_attention_pair_bias

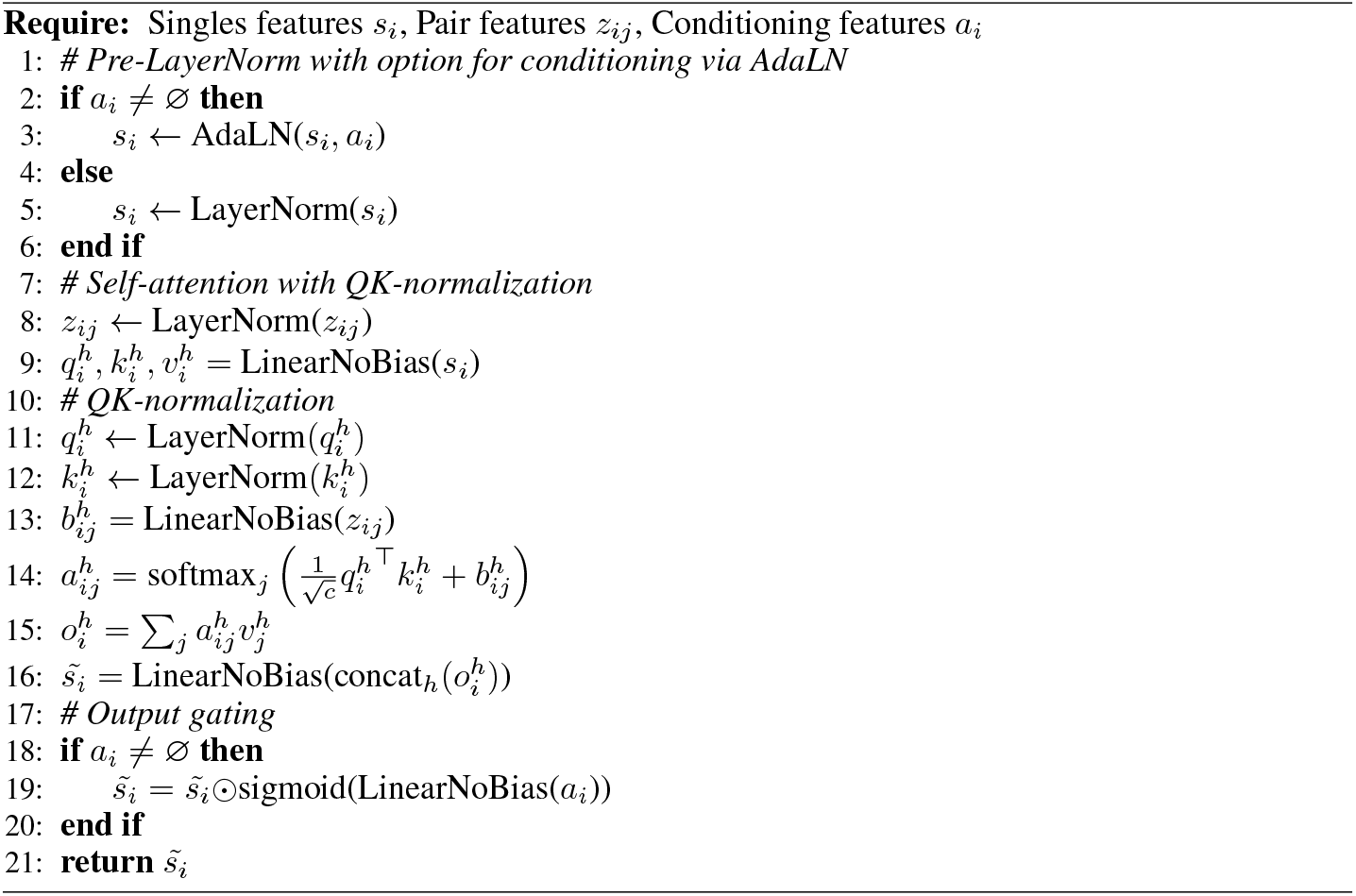

##### Algorithm 11

conditioned_transformer_pair_bias

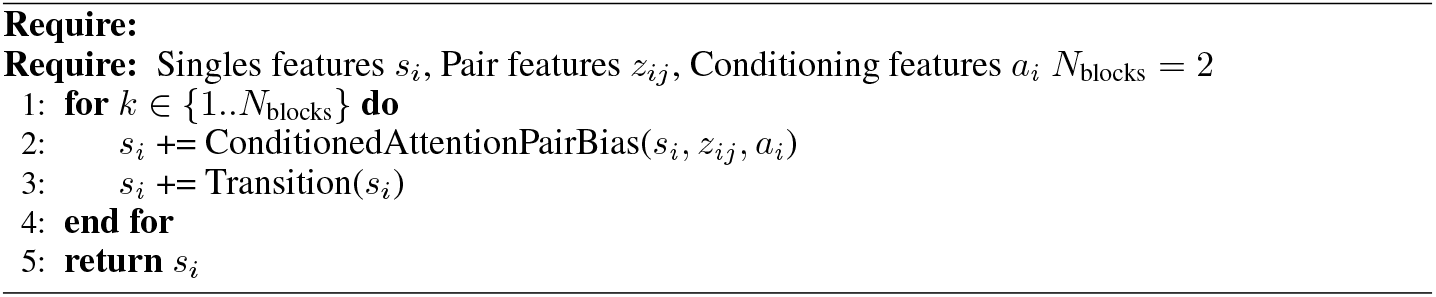

##### Algorithm 12

triangle_update

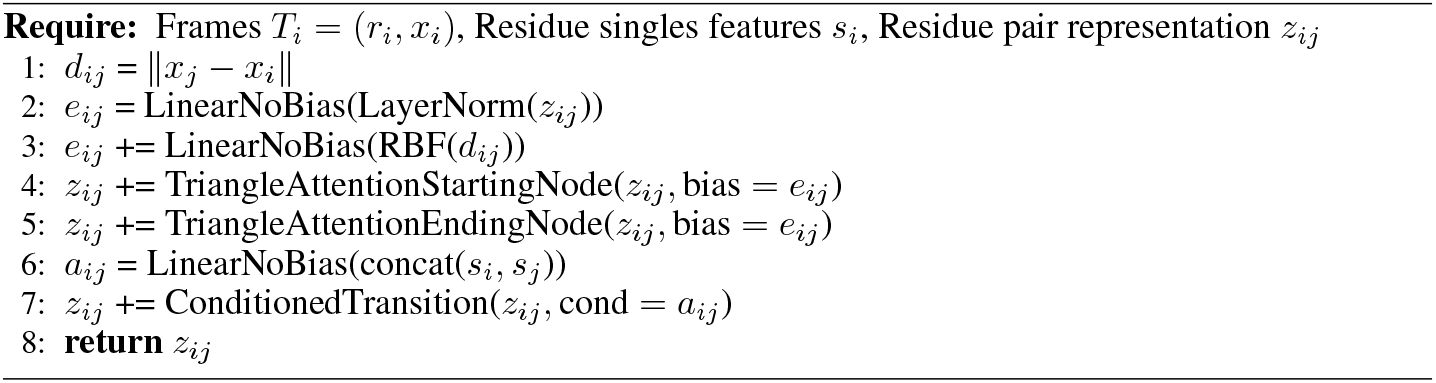

##### Algorithm 13

transition

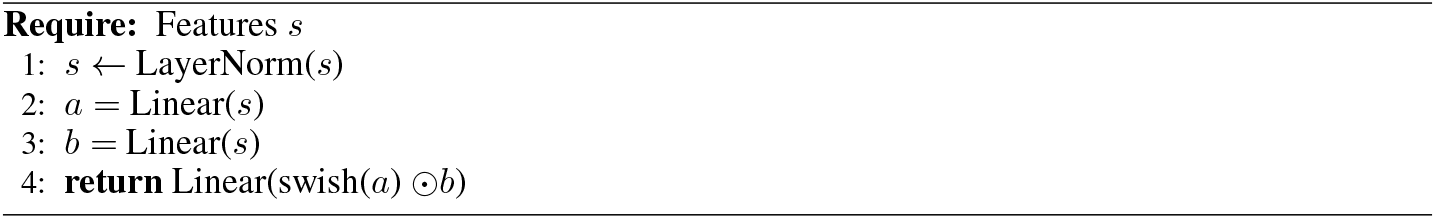

##### Algorithm 14

conditioned_transition

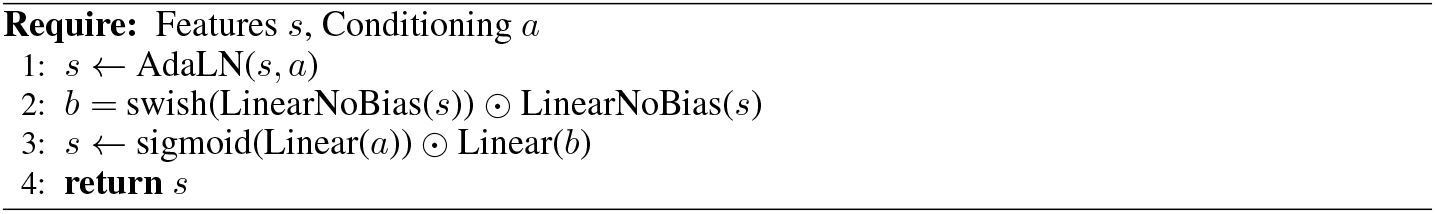

### F Sampling details

We describe our integrator for SDE sampling in Algorithm <u>15.</u> We use the following default parameters, but note that these can be tuned per user specification:

- Number of integration steps *T* = 400
- Noise scale parameter *γ*_*s*_ = 0.16
- Churn parameter *γ*_*c*_ = 0.4
- Translation Langevian schedule *g*_*x*_(*t*) = (1 − *t*)*/*(*t* + 0.1)^2^
- Rotation Langevian schedule *g*_*r*_(*t*) = (1 − *t*)*/*(*t* + 0.1)^2^
- Step scale *η* = 1.5
- Low noise threshold *T*_*ODE*_ = 0.99

For example, we find that changing the noise scale parameter *γ*_*s*_ allows us to tradeoff sample SSC and diversity (Figure F.1). We also compare against the classic SE(3) ODE sampler, which uses Euler integration with *T* = 400 to integrate the following ODEs:

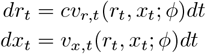

**Figure F.1:**
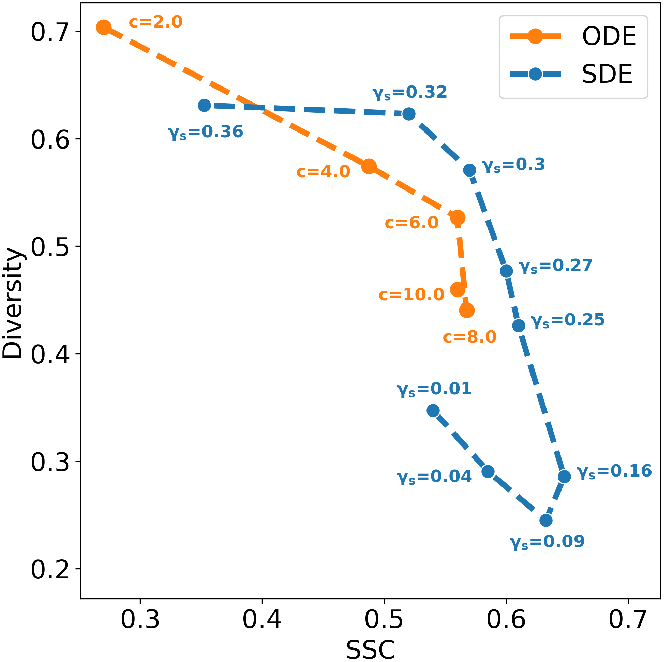
Tradeoff between diversity and SSC for unconditional monomer generation benchmark. We scan *γ*_*s*_ for the SDE (see Table F.1) and *c* for the ODE (see Table F.2). All other default parameters remain the same.

**Table F.1:** Raw values for *γ*_*s*_ scan for SDE sampling in Figure F.1.

| $\gamma_s$ | SSC | Diversity |
| --- | --- | --- |
| 0.01 | 0.54 | 0.35 |
| 0.04 | 0.59 | 0.29 |
| 0.09 | 0.63 | 0.25 |
| 0.16 | 0.65 | 0.29 |
| 0.25 | 0.61 | 0.43 |
| 0.27 | 0.60 | 0.48 |
| 0.30 | 0.57 | 0.57 |
| 0.32 | 0.52 | 0.62 |
| 0.36 | 0.35 | 0.63 |

**Table F.2:** Raw values for *c* scan for ODE sampling in Figure F.1.

| $c$ | SSC | Diversity |
| --- | --- | --- |
| 2 | 0.27 | 0.70 |
| 4 | 0.49 | 0.57 |
| 6 | 0.56 | 0.53 |
| 8 | 0.57 | 0.44 |
| 10 | 0.56 | 0.46 |

#### Algorithm 15

SDE sampling from ProteinZen

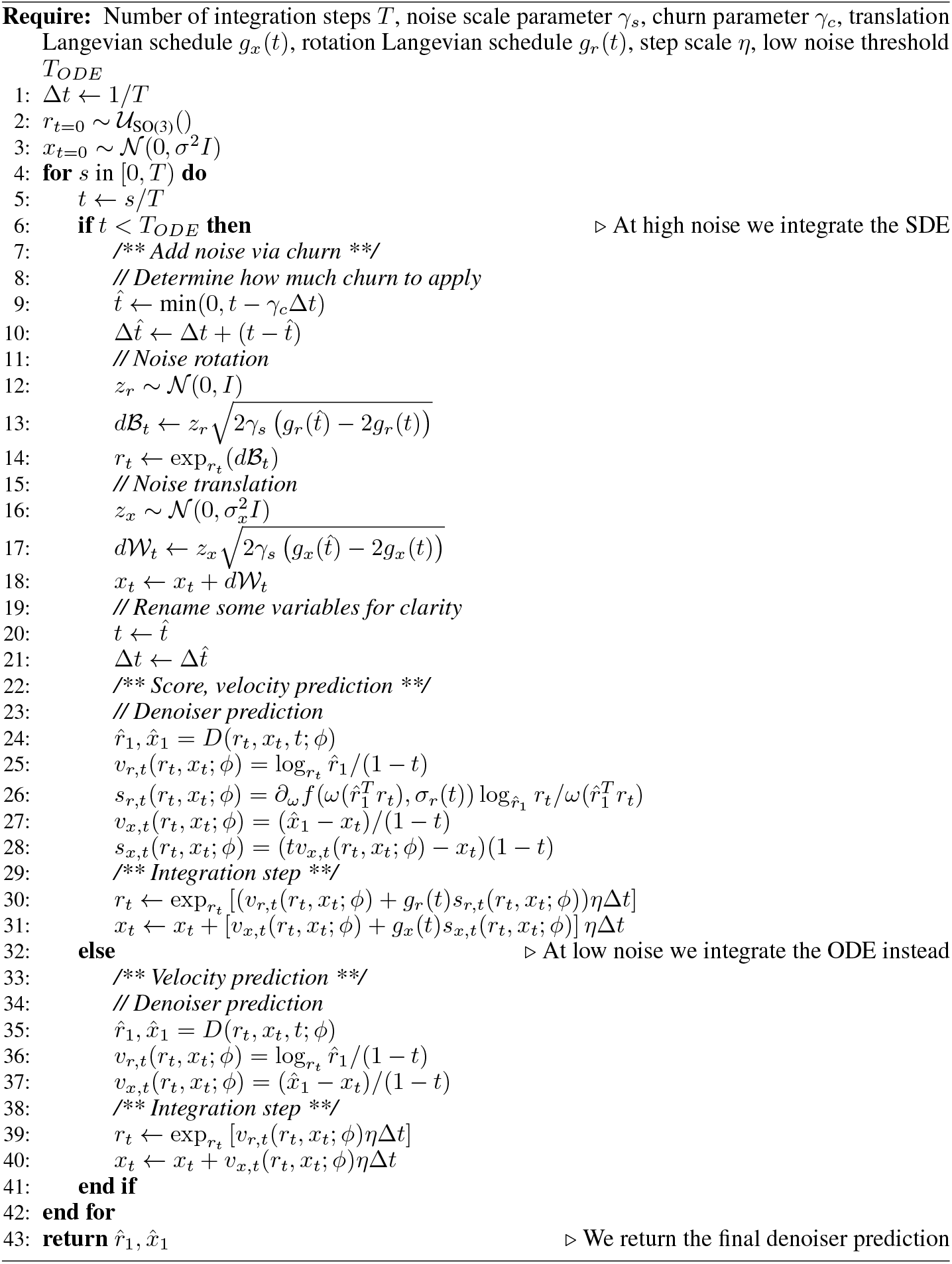

### G Unconditional monomer generation additional details

**Figure G.1:**
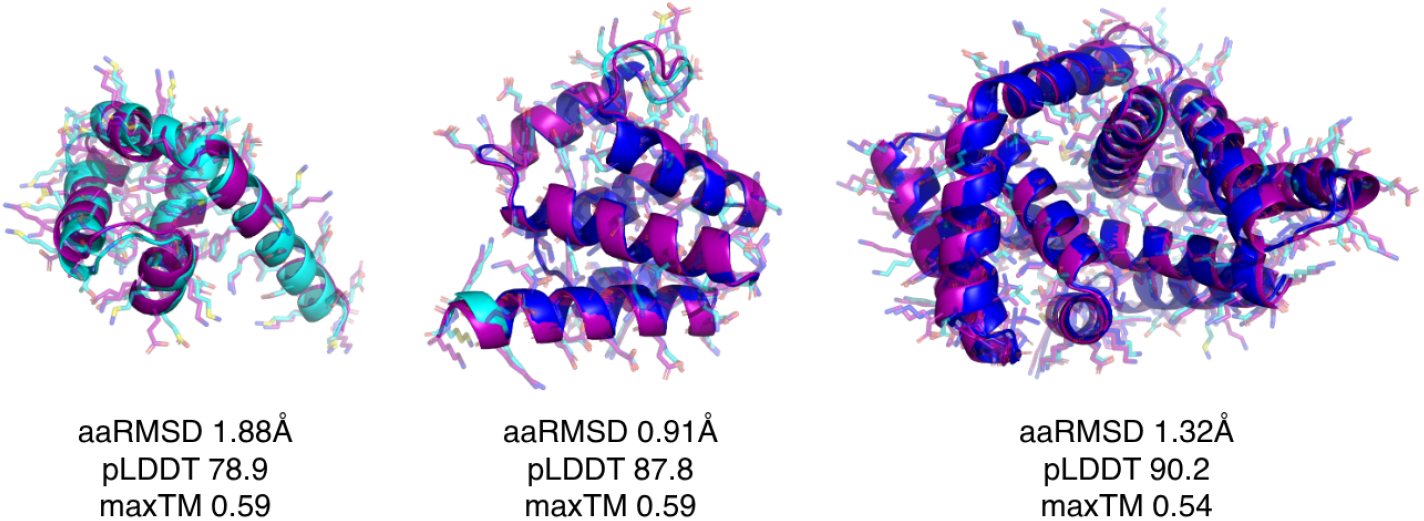
Samples from unconditional monomer generation with pdbTM < 0.6

### H Motif scaffolding additional details

#### H.1 Benchmark details

Task specifications for the Protpardelle motif scaffolding challenge are provided in Appendix Table H.1, where 200 designs are generated per task listed. Under indexed motif scaffolding, the model is given access to both motif order and spacing, whereas under unindexed motif scaffolding only the motif residues are provided.

**Table H.1:** Protpardelle motif scaffolding challenge specification.

| Task | Minimum length | Maximum Length | Task contig |
| --- | --- | --- | --- |
| 1BCF | 96 | 152 | 8-15/A92-99/16-30/A123-130/16-30/A47-54/16-30/A18-25/8-15 |
| 1PRW | 60 | 105 | 5-20/A16-35/10-25/A52-71/5-20 |
| 1QJG | 53 | 103 | 10-20/A38/15-30/A14/15-30/A99/10-20 |
| 1QJG_NATIVE | 115 | 135 | 10-20/A14/15-30/A38/50-70/A99/25-30 |
| 1YCR | 40 | 100 | 10-40/B19-27/10-40 |
| 2KL8 | 79 | 79 | A1-7/20/A28-79 |
| 3IXT | 50 | 75 | 10-40/P254-277/10-40 |
| 4JHW | 60 | 90 | 10-25/F196-212/15-30/F63-69/10-25 |
| 4ZYP | 30 | 50 | 10-40/A422-436/10-40 |
| 5AOU | 230 | 270 | 40-60/A1051/20-40/A2083/20-35/A2110/100-140 |
| 5AOU_QUAD | 230 | 270 | 40-60/A1051/20-40/A2083/20-35/A2110/60-80/A2180/40-60 |
| 5IUS | 57 | 142 | 0-30/A119-140/15-40/A63-82/0-30 |
| 5TPN | 50 | 75 | 10-40/A163-181/10-40 |
| 5TRV_SHORT | 56 | 56 | 0-35/A45-65/0-35 |
| 5TRV_MED | 86 | 86 | 0-65/A45-65/0-65 |
| 5TRV_LONG | 116 | 116 | 0-95/A45-65/0-95 |
| 5WN9 | 35 | 50 | 10-40/A170-189/10-40 |
| 5YUI | 50 | 100 | 5-30/A93-97/5-20/A118-120/10-35/A198-200/10-30 |
| 6E6R_SHORT | 48 | 48 | 0-35/A23-35/0-35 |
| 6E6R_MED | 78 | 78 | 0-65/A23-35/0-65 |
| 6E6R_LONG | 108 | 108 | 0-95/A23-35/0-95 |
| 6VW1 | 62 | 83 | 20-30/A24-42/4-10/A64-82/0-5 |
| 7K4V | 280 | 320 | 40-50/A44/3-8/A50/70-85/A127/150-200 |
| 7MRX_60 | 60 | 60 | 0-38/B25-46/0-38 |
| 7MRX_85 | 85 | 85 | 0-63/B25-46/0-63 |
| 7MRX_128 | 128 | 128 | 0-122/B25-46/0-122 |

We provide the raw task success metrics used to create Figure 3 in Table H.2 and Table H.3, where values are computed from samples generated using the published implementations:

- La-Proteina: commit `8f74f7990ab9e1ac1169d8d9921b2e09425701e9`
- Protpardelle-1c: commit `9c404865167fc08fabbf07086381d3248d787af1`
- RFDiffusion: commit `820bfdfaded8c260b962dc40a3171eae316b6ce0`

We note that benchmark values for comparison models are slightly lower across the board when compared to reference literature [13][16], but that all designs were evaluated using a common evaluation pipeline so benchmark values are directly comparable.

**Table H.2:** Total and unique number of successes per task on one-shot Protpardelle motif scaffolding benchmark.

| Task | Protpardelle-1c cc91 |  | La-Proteina (indexed) |  | La-Proteina (unindexed) |  | ProteinZen (indexed) |  | ProteinZen (unindexed) |  | # segments |
| --- | --- | --- | --- | --- | --- | --- | --- | --- | --- | --- | --- |
|  | All | Unique | All | Unique | All | Unique | All | Unique | All | Unique |  |
| 1YCR | 50 | 22 | 118 | 45 | 105 | 67 | 125 | 67 | 132 | 86 | 1 |
| 3IXT | 2 | 2 | 39 | 4 | 53 | 6 | 31 | 12 | 23 | 9 | 1 |
| 4ZYP | 4 | 2 | 3 | 1 | 29 | 3 | 26 | 8 | 18 | 7 | 1 |
| 5TPN | 17 | 6 | 33 | 6 | 28 | 9 | 32 | 13 | 38 | 13 | 1 |
| 5TRV_short | 0 | 0 | 1 | 1 | 0 | 0 | 1 | 1 | 1 | 1 | 1 |
| 5TRV_med | 0 | 0 | 27 | 10 | 31 | 7 | 6 | 4 | 10 | 8 | 1 |
| 5TRV_long | 0 | 0 | 48 | 18 | 59 | 28 | 15 | 12 | 12 | 11 | 1 |
| 5WN9 | 0 | 0 | 0 | 0 | 0 | 0 | 6 | 4 | 2 | 2 | 1 |
| 6E6R_short | 5 | 3 | 23 | 15 | 46 | 20 | 22 | 16 | 20 | 17 | 1 |
| 6E6R_med | 19 | 11 | 89 | 43 | 77 | 50 | 62 | 50 | 70 | 54 | 1 |
| 6E6R_long | 11 | 11 | 80 | 46 | 93 | 69 | 71 | 58 | 69 | 59 | 1 |
| 7MRX_60 | 8 | 3 | 36 | 3 | 38 | 3 | 50 | 3 | 47 | 5 | 1 |
| 7MRX_85 | 13 | 5 | 23 | 3 | 64 | 6 | 54 | 9 | 47 | 5 | 1 |
| 7MRX_128 | 2 | 2 | 9 | 5 | 64 | 21 | 16 | 9 | 17 | 6 | 1 |
| 1PRW | 148 | 13 | 167 | 18 | 115 | 9 | 139 | 36 | 50 | 17 | 2 |
| 2KL8 | 78 | 1 | 63 | 1 | 27 | 1 | 11 | 1 | 13 | 1 | 2 |
| 4JHW | 0 | 0 | 0 | 0 | 0 | 0 | 0 | 0 | 0 | 0 | 2 |
| 5IUS | 0 | 0 | 1 | 1 | 0 | 0 | 3 | 2 | 0 | 0 | 2 |
| 6VW1 | 12 | 1 | 28 | 1 | 37 | 1 | 45 | 2 | 56 | 9 | 2 |
| 1QJG | 4 | 4 | 28 | 14 | 49 | 27 | 41 | 24 | 47 | 38 | 3 |
| 1QJG_NAT | 10 | 8 | 39 | 15 | 83 | 51 | 39 | 22 | 58 | 43 | 3 |
| 5AOU | 38 | 4 | 151 | 8 | 20 | 16 | 63 | 14 | 4 | 2 | 3 |
| 5YUI | 0 | 0 | 3 | 2 | 8 | 8 | 1 | 1 | 1 | 1 | 3 |
| 7K4V | 4 | 2 | 40 | 8 | 23 | 23 | 4 | 1 | 1 | 1 | 3 |
| 1BCF | 195 | 6 | 144 | 7 | 90 | 6 | 175 | 9 | 138 | 22 | 4 |
| 5AOU_QUAD | 14 | 3 | 81 | 5 | 50 | 25 | 68 | 11 | 4 | 2 | 4 |
| **Total** | **634** | **109** | **1274** | **280** | **1189** | **456** | **1106** | **389** | **878** | **419** |  |

**Table H.3:** Total and unique number of successes per task on Protpardelle motif scaffolding benchmark with sequence redesign by ProteinMPNN.

| Task | RFDiffusion |  | Protpardelle-1c cc91 |  | La-Proteina (indexed) |  | La-Proteina (unindexed) |  | ProteinZen (indexed) |  | ProteinZen (unindexed) |  | # segments |
| --- | --- | --- | --- | --- | --- | --- | --- | --- | --- | --- | --- | --- | --- |
|  | All | Unique | All | Unique | All | Unique | All | Unique | All | Unique | All | Unique |  |
| 1YCR | 171 | 75 | 134 | 55 | 175 | 70 | 182 | 107 | 198 | 112 | 200 | 121 | 1 |
| 3IXT | 145 | 32 | 15 | 4 | 81 | 12 | 158 | 23 | 158 | 58 | 153 | 49 | 1 |
| 4ZYP | 124 | 23 | 30 | 9 | 85 | 15 | 153 | 21 | 145 | 44 | 153 | 56 | 1 |
| 5TPN | 168 | 42 | 56 | 30 | 79 | 19 | 92 | 30 | 171 | 60 | 157 | 84 | 1 |
| 5TRV_short | 0 | 0 | 0 | 0 | 8 | 3 | 10 | 2 | 171 | 41 | 174 | 45 | 1 |
| 5TRV_med | 4 | 4 | 0 | 0 | 85 | 19 | 112 | 32 | 152 | 56 | 183 | 65 | 1 |
| 5TRV_long | 24 | 18 | 0 | 0 | 126 | 33 | 161 | 57 | 164 | 78 | 187 | 90 | 1 |
| 5WN9 | 1 | 1 | 0 | 0 | 2 | 2 | 2 | 2 | 40 | 21 | 36 | 21 | 1 |
| 6E6R_short | 79 | 33 | 53 | 29 | 87 | 34 | 124 | 53 | 163 | 105 | 168 | 106 | 1 |
| 6E6R_med | 160 | 60 | 69 | 38 | 153 | 79 | 171 | 110 | 174 | 126 | 184 | 127 | 1 |
| 6E6R_long | 149 | 72 | 53 | 45 | 163 | 92 | 172 | 137 | 183 | 141 | 190 | 137 | 1 |
| 7MRX_60 | 41 | 9 | 55 | 12 | 124 | 8 | 129 | 4 | 123 | 18 | 121 | 20 | 1 |
| 7MRX_85 | 27 | 13 | 59 | 15 | 120 | 15 | 173 | 10 | 133 | 24 | 135 | 25 | 1 |
| 7MRX_128 | 15 | 11 | 16 | 14 | 68 | 24 | 120 | 43 | 114 | 30 | 113 | 34 | 1 |
| 1PRW | 199 | 15 | 195 | 21 | 199 | 16 | 176 | 12 | 194 | 30 | 88 | 23 | 2 |
| 2KL8 | 180 | 1 | 164 | 1 | 185 | 1 | 19 | 1 | 178 | 1 | 54 | 1 | 2 |
| 4JHW | 0 | 0 | 1 | 1 | 5 | 4 | 1 | 1 | 24 | 16 | 15 | 11 | 2 |
| 5IUS | 14 | 2 | 2 | 2 | 8 | 2 | 0 | 0 | 133 | 22 | 67 | 26 | 2 |
| 6VW1 | 149 | 7 | 71 | 7 | 122 | 2 | 156 | 10 | 181 | 11 | 187 | 33 | 2 |
| 1QJG | 102 | 62 | 42 | 29 | 123 | 40 | 139 | 61 | 155 | 73 | 171 | 130 | 3 |
| 1QJG_NAT | 113 | 56 | 59 | 36 | 139 | 41 | 176 | 114 | 162 | 95 | 168 | 127 | 3 |
| 5AOU | 44 | 34 | 98 | 9 | 197 | 10 | 61 | 53 | 131 | 42 | 36 | 8 | 3 |
| 5YUI | 2 | 2 | 8 | 5 | 16 | 12 | 43 | 22 | 78 | 63 | 74 | 71 | 3 |
| 7K4V | 40 | 36 | 16 | 3 | 125 | 21 | 68 | 66 | 51 | 17 | 54 | 27 | 3 |
| 1BCF | 200 | 10 | 200 | 6 | 196 | 10 | 199 | 13 | 190 | 13 | 197 | 34 | 4 |
| 5AOU_QUAD | 29 | 28 | 62 | 7 | 171 | 9 | 117 | 59 | 161 | 22 | 26 | 20 | 4 |
| **Total** | **2180** | **646** | **1458** | **378** | **2842** | **593** | **2914** | **1043** | **3727** | **1319** | **3291** | **1491** |  |

#### H.2 Scaffold characteristics

We provide example task successes for indexed motif scaffolding in Figure H.1 and unindexed motif scaffolding in Figure H.2. We also plot RMSD metrics for indexed motif scaffolding in Figures H.3, H.4 and for unindexed motif scaffolding in Figures H.5, H.6. Similar to under unconditional monomer generation, ProteinZen appears to have a preference for generating helical elements in scaffolds, but is both capable of scaffolding motifs with *β*-strands and also generating scaffolds with *β*-sheets in them.

**Figure H.1:**
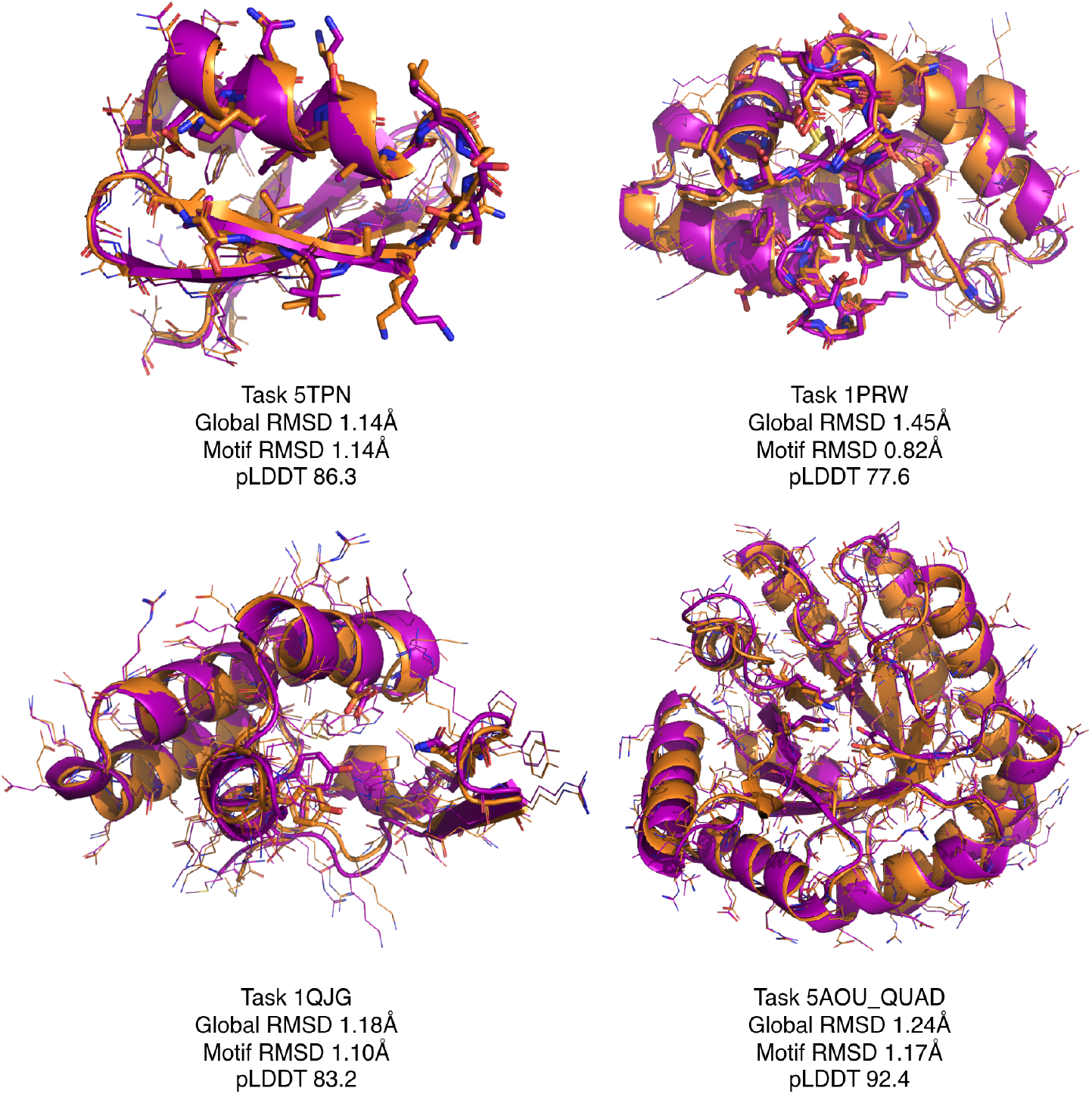
Examples of indexed motif scaffolding task successes. Purple structures are the design structure, and orange structures are the corresponding ESMFold prediction. Atomic detail is shown in wireframe, and motif residues are displayed using sticks.

**Figure H.2:**
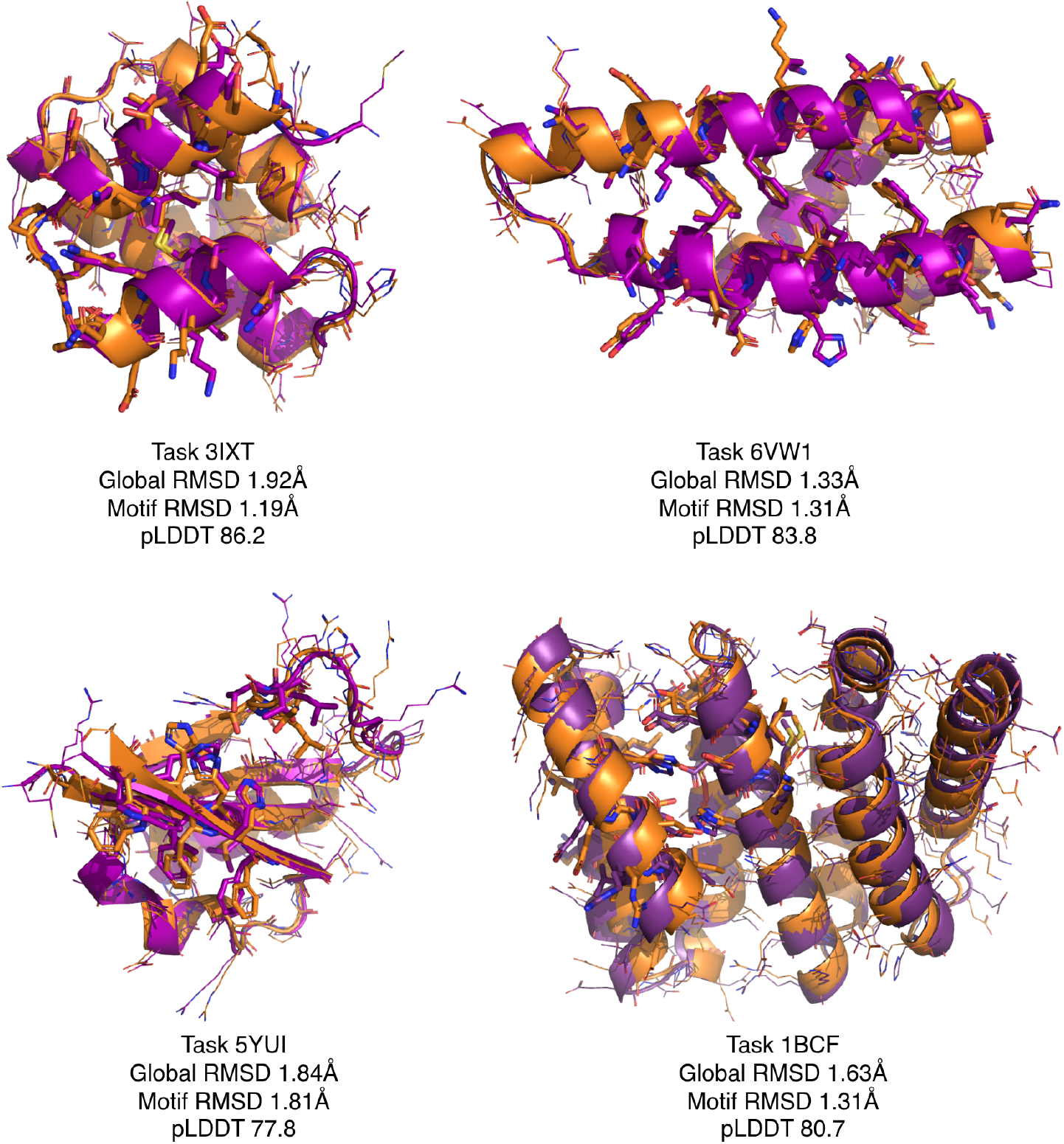
Examples of unindexed motif scaffolding task successes. Purple structures are the design structure, and orange structures are the corresponding ESMFold prediction. Atomic detail is shown in wireframe, and motif residues are displayed using sticks.

**Figure H.3:**
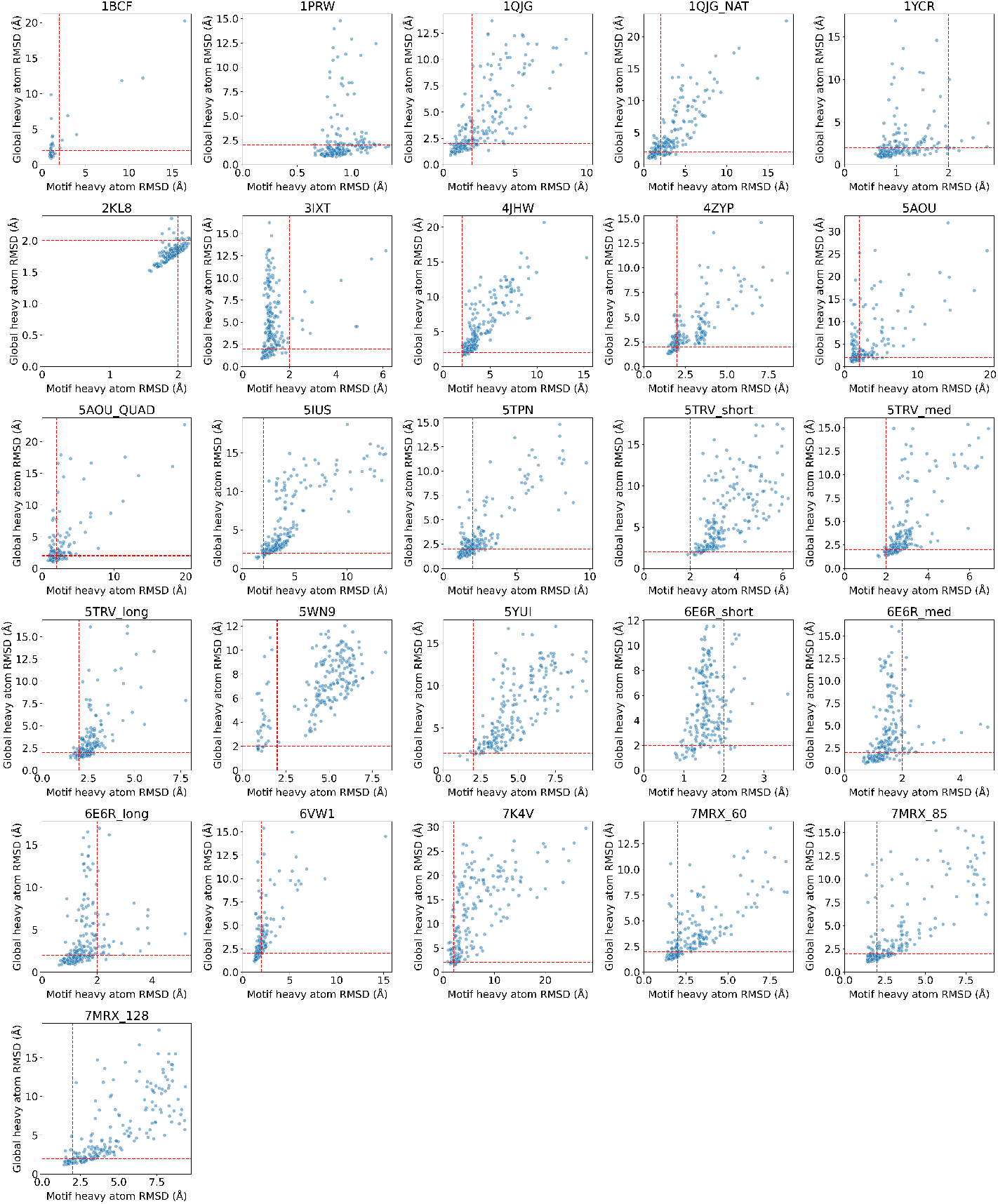
Motif heavy atom RMSD vs global heavy atom RMSD for indexed motif scaffolding designs. Red lines mark motif heavy atom RMSD < 2Å and global heavy atom RMSD < 2Å.

**Figure H.4:**
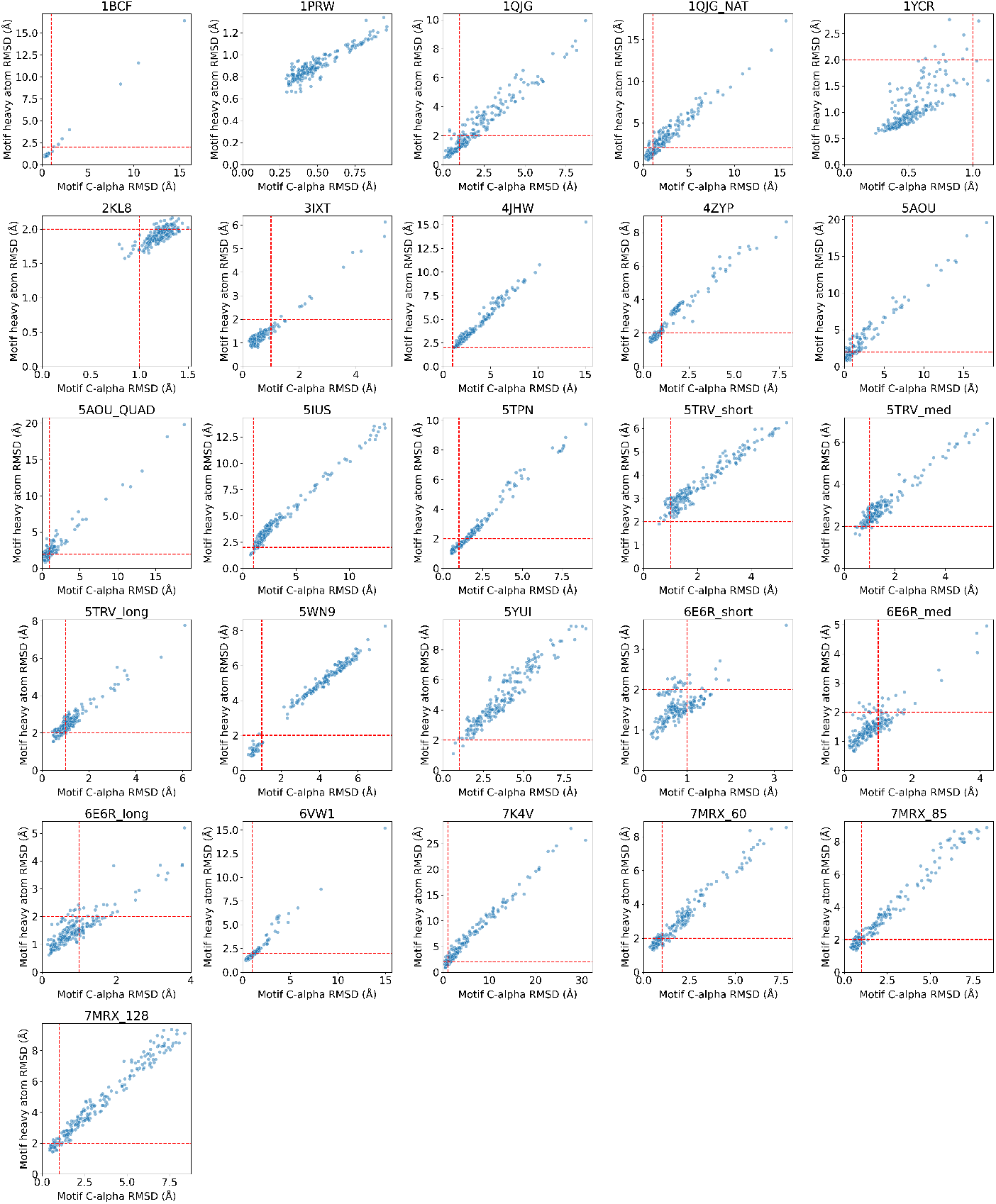
Motif C*α* RMSD vs motif heavy atom RMSD for indexed motif scaffolding designs. Red lines mark motif C*α* RMSD < 1Å and motif heavy atom RMSD < 2Å.

**Figure H.5:**
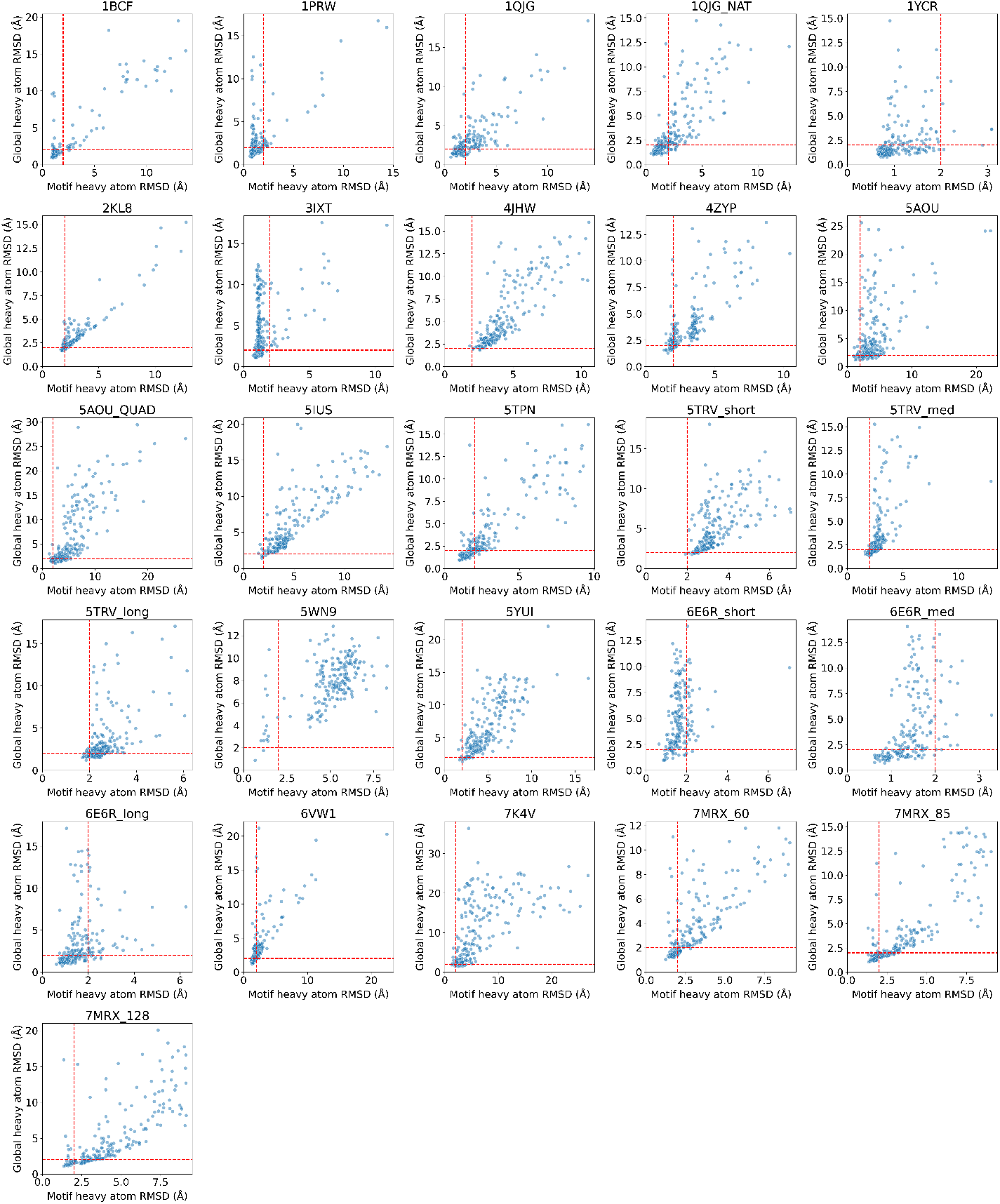
Motif heavy atom RMSD vs global heavy atom RMSD for unindexed motif scaffolding designs. Red lines mark motif heavy atom RMSD < 2Å and global heavy atom RMSD < 2Å.

**Figure H.6:**
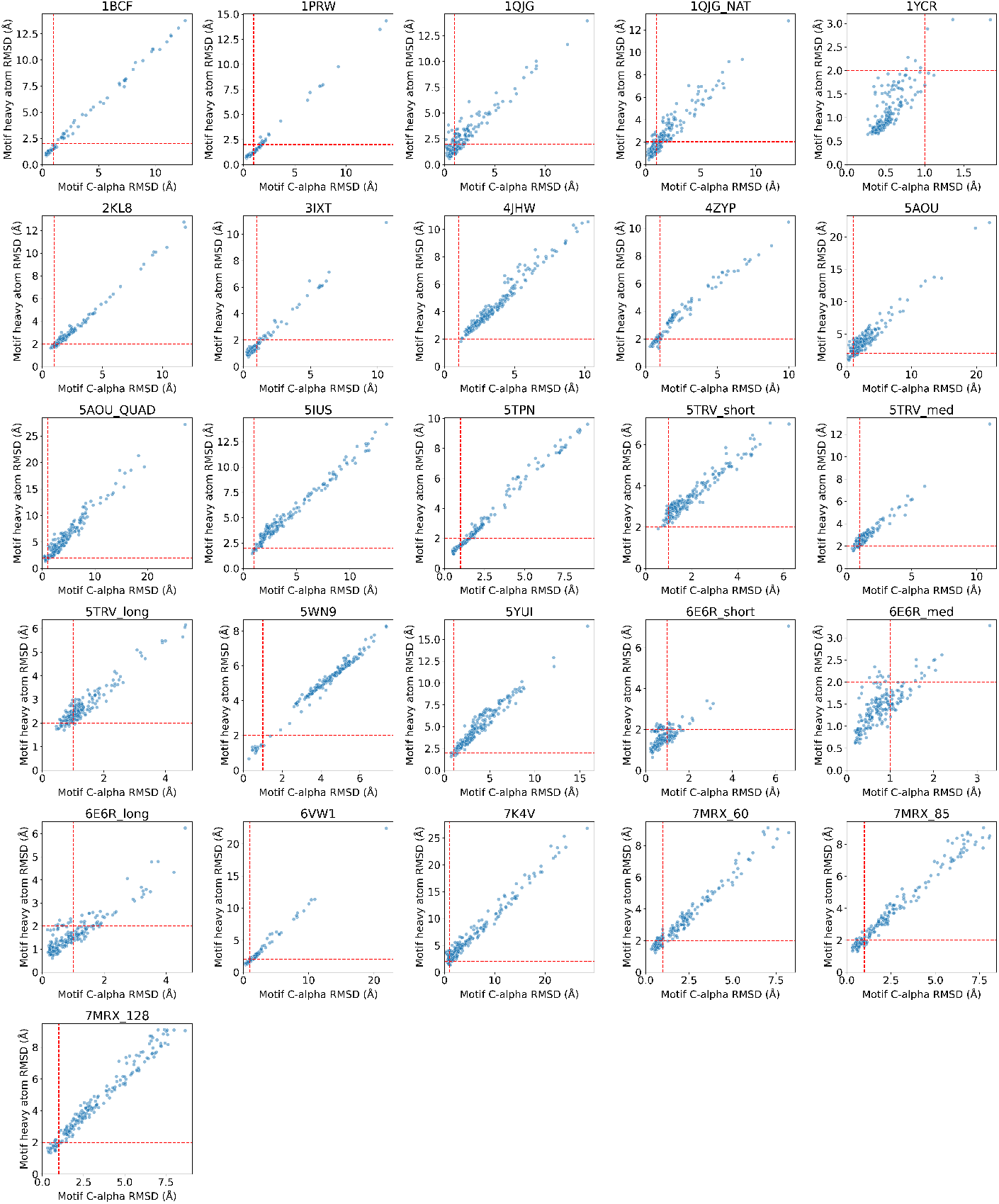
Motif C*α* RMSD vs motif heavy atom RMSD for unindexed motif scaffolding designs. Red lines mark motif C*α* RMSD < 1Å and motif heavy atom RMSD < 2Å.

## Footnotes

1 Code is available at https://github.com/alexjli/proteinzen.

